# The kinetochore corona orchestrates chromosome congression through transient microtubule interactions

**DOI:** 10.1101/2025.09.10.675486

**Authors:** Christopher E. Miles, Fioranna Renda, Irina Tikhonenko, Angus Alfieri, Alex Mogilner, Alexey Khodjakov

## Abstract

For proper segregation of chromosomes and successful cytokinesis, chromosomes must first ‘congress’ - gather in a tight plate near the spindle equator. Molecular mechanism(s) of congression are not fully understood. Here we combine live cell microscopy, perturbations of microtubule motor activities, correlative light/electron microscopy, and computational modeling, to quantitatively characterize the early-prometaphase movements that bring the scattered chromosomes to the equator in human RPE1 cells. We find that the early-prometaphase movements are predominantly directed toward the spindle center and not the spindle poles. Centromere velocity of the centripetal movements is not constant with centromeres moving faster at larger distances from the spindle center. We also detect that numerous short microtubules appear at kinetochores at the earliest stages of spindle assembly and prior to chromosome congression. Computational modeling reveals that a mechanism based on brief, stochastic, minus-end directed interactions between the short microtubules protruding from the kinetochores and long appropriately curved microtubules within the spindle accurately predicts the observed distance-velocity function. Further, the model predicts that insufficient numbers of microtubules protruding from the kinetochores decreases the velocity and randomizes directionality of congression movements. These predictions match changes in the chromosome behavior observed in cells with suppressed nucleation of microtubules at the kinetochore corona (RPE1 Rod^Δ/Δ^). In contrast, predictions of computational models based on continuous pulling forces at kinetochores differ significantly from the experimental observations. Together, live-cell observations and modeling reveal a novel mechanism that enables the efficient and synchronized arrival of chromosomes to the spindle equator.

**Significance Statement:** For equal segregation into daughter cells, chromosomes, scattered in a large volume at the onset of cell division, must congress into a narrow plate near the equator of the mitotic spindle. Molecular mechanisms of congression remain obscure. Here we use live-cell microscopy and structural analyses to quantitatively characterize chromosome behavior during congression in human cells. From these quantifications we derive a computational model that accurately predicts directionality and velocity of chromosome movements and suggests that the force for these movements arises from stochastic, transient, minus-end directed interactions between short microtubules protruding from the kinetochores and long appropriately shaped microtubules within the spindle. The model also accurately predicts changes in chromosome behavior in cells with functionally deficient kinetochores.

## Introduction

Equal segregation of chromosomes into the two daughter cells during mitosis is enacted by the ‘mitotic spindle’, a self-assembling macromolecular machine comprising thousands of microtubules (MTs) (1–3). The two sister chromatids move to the opposite poles of the spindle by forces acting along MTs attached to the ‘kinetochores’, two macromolecular complexes residing on the opposite sides of chromosome’s ‘centromere’. MTs attached to kinetochores form bundles of uniform polarity (‘K-fibers’) with MT plus ends residing at the kinetochore (4). Two mechanisms of K-fiber formation have been identified. In one, a kinetochore attaches to MTs produced by a spindle pole which instantly establishes a direct connection with the pole (5). Alternatively, kinetochores grab MTs nucleated in their immediate proximity and the connection to the pole is mediated by interactions between the distal (minus) ends of short MTs protruding from the kinetochore (skMTs) and other MTs within the spindle (1, 6–8).

Formation of simultaneous connections between sister kinetochores and the opposite spindle poles (termed ‘amphitelic attachment’) is most efficient when centromeres reside near the spindle equator equidistant to the poles (9). In contrast, near a spindle pole establishing connection to the distal pole is impeded. Indeed, conditions that promote accumulation of chromosomes near spindle poles (termed ‘monoorientation’) decrease the fidelity of chromosome segregation and adversely affect the progeny (10). Thus, a major goal of spindle assembly is to rapidly gather the initially scattered chromosomes in a tight group at the spindle equator. Molecular mechanism(s) that drive this gathering (termed ‘chromosome congression’) are not well understood (11). Here, we combine quantitative analysis of chromosome behavior during early stages of spindle assembly (prometaphase), correlative light/electron microscopy (CLEM) reconstructions, and computational modeling to reveal these mechanisms. Our analyses suggest that direct kinetochore attachment to MTs produced by spindle poles are infrequent and the rapid chromosome congression during early prometaphase is driven primarily by highly dynamic interactions between skMTs and long, properly shaped, anti-parallel MTs within the spindle. The force generated by these interactions is directed toward the minus ends of spindle MTs, which suggests involvement of cytoplasmic dynein. Importantly, our computational model accurately predicts measurable changes in early-prometaphase chromosome behavior, including attenuated velocity of centromere movements and the rate of monoorientation corrections, that occur in cells with a lower number of skMTs. This represents a significant step towards a truly quantitative and predictive computational model of mitotic spindle assembly in human cells.

## Results

### Early-prometaphase decrease in monooriented chromosomes depends on the kinetochore corona but not on the plus-end directed motor activity at the kinetochore

In human somatic cells, the mitotic spindle principally comprises two asters of MTs that emanate from the spindle poles and overlap in the middle (Fig. 1A and SI Appendix, Fig. S1A). In this architecture, chromosomes residing near the spindle equator (termed ‘bioriented’) attach simultaneously to MTs emanating from both spindle poles (termed ‘amphitelic attachment’), which is necessary for proper segregation. In contrast, formation of amphitelic attachments on ‘monooriented’ chromosomes, near a spindle pole, is improbable due to the lower number of MTs from the distal pole (Fig. 1A). These chromosomes need to reposition closer to the equator (termed ‘congression’, Fig. 1B).

**Fig. 1.**
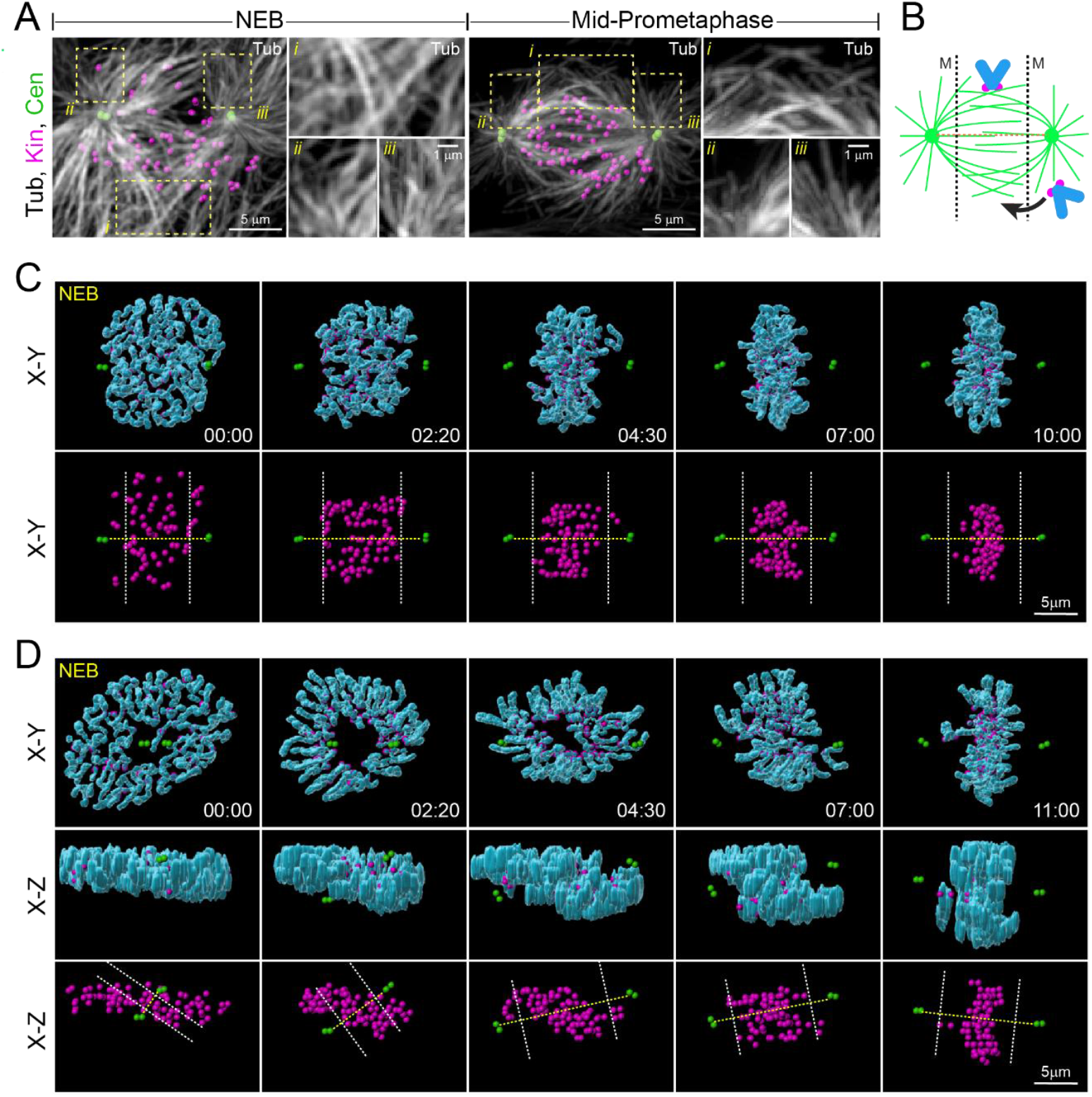
Formation of metaphase plate in RPE1 cells with steadily horizontal or gradually rotating spindle. **(A)** Distribution of MTs and centromeres at nuclear envelope breakdown (NEB) and during mid-prometaphase. Maximum-intensity projections of the entire cell. Insets show boxed areas at a higher magnification. *i)* overlapping MTs near the spindle equator, *ii)* and *iii)* radial microtubule arrays near spindle poles. MTs (white), centrioles (green) and kinetochores (magenta) are shown. **(B)** Cartoon of the spindle during prometaphase. Chromosome near the equator encounter MTs from both poles while near spindle poles MTs emanating from the distal spindle pole are rare. Dashed lines denote parts of the spindle where chromosomes are ‘monooriented’. **(C-D)** Changes in spatial distribution of chromosomes during prometaphase in cells with steadily horizontal (C) or rotating (D) spindle. The volumes are aligned at each time point to stabilize XYZ position of the spindle center and spindle orientation in XY (but not in XZ) projections. Centrosomes are on the opposite sides of the nucleus along the longer (C) or shorter (D) axis of the discoid nucleus at NEB. Notice that chromosomes rapidly void monoorientation zones (outside of dashed white lines). Dashed yellow lines denote the spindle axis. Chromosome arms are segmented and surface-rendered (blue). Balls mark positions of kinetochores (magenta) and centrioles (green). Timestamps are in minutes : seconds.

The spindle assembly initiates when the nuclear envelope breakdown (NEB) enables direct contacts between the chromosomes and MTs. At NEB, chromosomes are uniformly scattered within the nuclear volume shaped as a discoid with the shorter axis orthogonal and longer axis parallel to the substrate. Two centrosomes, whose MT nucleating activities define the spindle poles, reside consistently on the opposite sides of the nucleus within invaginations of the nuclear envelope (SI Appendix, Fig. S1A, NEB). In cells where centrosomes are separated at NEB along the longer nuclear axis, the spindle axis remains roughly parallel to the substrate and the length of the spindle is relatively constant throughout mitosis. Although infrequent (~25% of RPE1 (12, 13)), these cells offer a stable viewpoint on the chromosome behavior throughout prometaphase (SI Appendix, Fig. S1B). Conventional microscopy reveals that the separation between the centrosomes and chromosomes increases progressively after NEB (Fig. 1C). In less than three minutes, chromosomes clear the monoorientation zones near spindle poles (Fig. 1C, 2:20) and only occasionally individual chromosomes transiently approach the poles at later times (Fig. 1C, 4:30). Kinetochores rapidly gather from a scattered cloud into a tight equatorial plate (Fig. 1C).

In most cells (~75% in RPE1), the centrosomes separate along the shorter nuclear axis so that the spindle initially is nearly orthogonal to the substrate but subsequently rotates to nearly parallel to the substrate by mid-late prometaphase (9, 12, 13). In these cells, chromosome movements within the spindle are obscured by the constantly changing viewpoint (SI Appendix, Fig. S1C). While chromosomes often appear near poles in the conventional projections onto XY-plane (SI Appendix, Fig. S1C), 3-D visualization reveals that the distance between chromosomes and the poles progressively increases (Fig. 1D) akin in cells with steadily horizontal spindle orientation.

To uncouple chromosome behavior within the spindle from changes in the spindle orientation within the cell, we express positions of chromosomes’ centromeres (defined here as the midpoint between sister kinetochores) in a cylindrical coordinate system defined by the distance to the spindle axis (ρ), distance to the spindle equator (Z), and angle between the horizon and the vector from the spindle center to the centromere (θ) (SI Appendix, Fig. S2A). This approach allows us to separately assess the axial (parallel to the spindle axis) and equatorial (parallel to the equatorial plane) components of centromere movements. Further, by normalizing Z values to the spindle length, we can directly compare centromere behavior within the monoorientation (<1/3 of the half-spindle length from a pole) and biorientation zones (SI Appendix, Fig. S2A).

Due to the initial position of centrosomes within invaginations of the nuclear envelope, ~40% of centromeres in RPE1 cells (358 of 876 tracked in 19 cells) are monooriented at NEB. Indeed, many centromeres are farther from the equator than the centrosomes at this stage (Fig. 2A). The number of the inherently monooriented “polar” centromeres decreases rapidly from ~20 to ~2 per cell in the first 400 s of prometaphase (Fig. 2B, RPE1) and the correction involves movement of centromeres closer to the equator (Fig. 2C, RPE1). In contrast, inherently “equatorial” centromeres rarely become monooriented later in prometaphase (Fig. 2A and B, RPE1) and their mean distance to the equator remains relatively constant (Fig. 2C, RPE1). To test whether the changes in distribution of polar centromeres requires a motor activity, we use 20-nM GSK923295 to inhibit CenpE, the only plus-end directed kinesin at the kinetochore (14). CenpE inhibition does not affect the decrease in monooriented chromosomes during the first 400 seconds of prometaphase (Fig. 2B, RPE1 CenpE-inh). Later, the number of monooriented centromeres increases in CenpE-inhibited RPE1 as some already-congressed chromosomes reposition closer to the spindle poles. Both “polar” and “equatorial” centromeres contribute to this process at the same rate (Fig. 2B and C, RPE1 CenpE-inh). Thus, while congression of the polar centromeres does not require a plus-end directed motor activity, CenpE appears to help maintaining equatorial positions of already congressed chromosomes later in prometaphase.

**Fig. 2.**
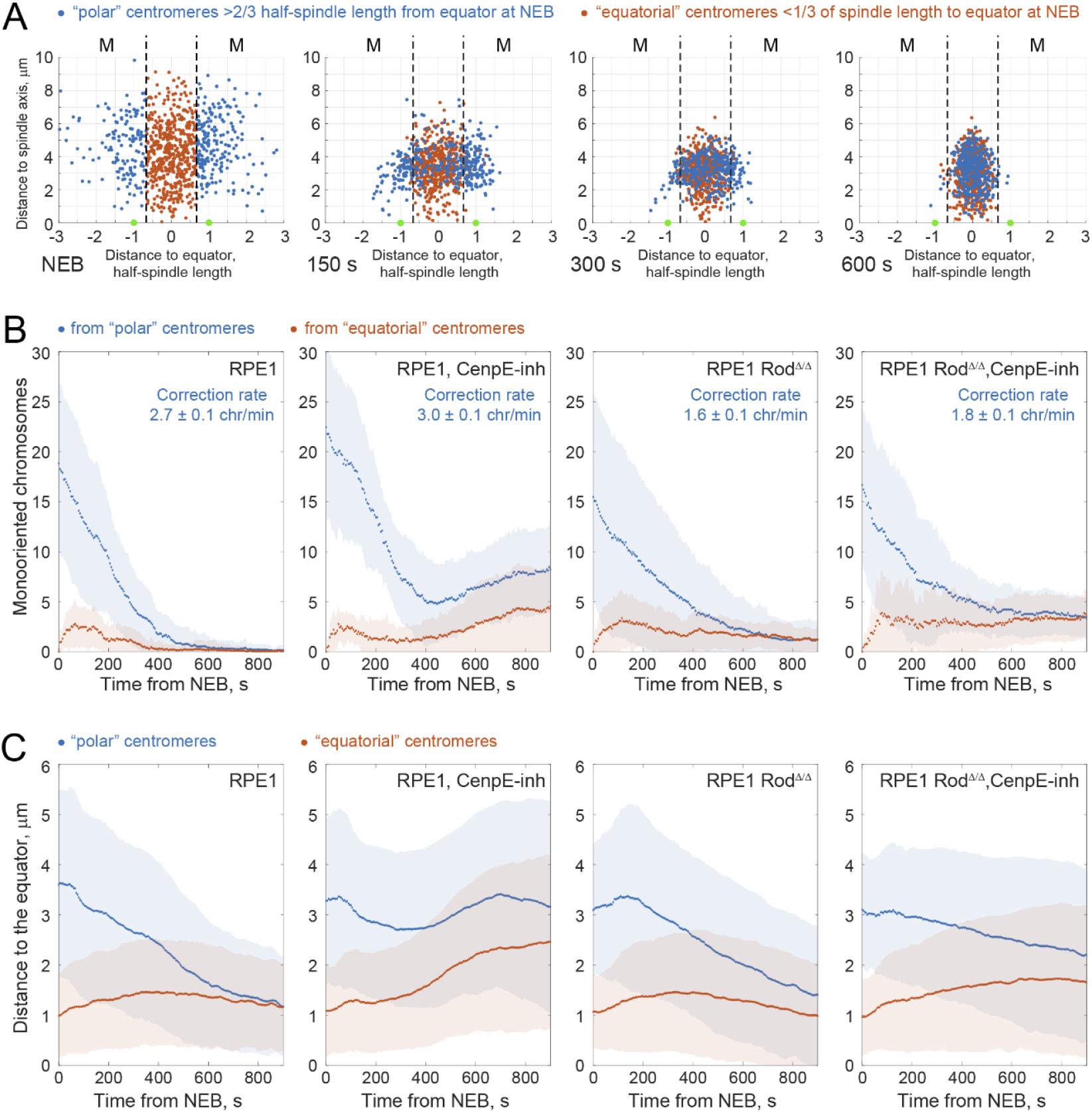
Correction of chromosome monoorientation depends on the kinetochore corona but not on a plus-end directed motor activity. **(A)** Z-ρ distribution of centromeres at various stages of spindle assembly in RPE1 cells (876 centromeres in 19 cells). “Polar” centromeres, that resided in the monoorientation zones (M, outside of black dashed line) at NEB, gradually relocate closer to the equator, while “equatorial” centromeres positioned near the equator at NEB rarely become monooriented. **(B)** Number of monooriented chromosomes (mean and SD per cell) from the “polar” (blue) and “equatorial” (orange) centromeres. “CenpE-inh” cells were treated with 20-nM GSK923295. Rates of monoorientation are the slope of linear regression calculated for the period from 5 to 60 seconds after NEB. **(C)** Mean (with STD) distance from the equator to “polar” (blue) and “equatorial” (orange) centromeres during prometaphase.

Monooriented chromosomes are known to become numerous when kinetochores lack the ‘fibrous corona’, an outer-kinetochore complex involved in capturing spindle MTs as well as for nucleating non-centrosomal MTs near kinetochores (15–17). To assess monoorientation dynamics in the absence of the corona we use RPE1 Rod^Δ/Δ^ cells (9, 18). In these cells, the number of polar chromosomes decreases at an ~60% slower rate than in the wild-type RPE1, irrespective of CenpE activity (Fig. 2B, Rod^Δ/Δ^). Further, congression of the initially polar centromeres is significantly delayed in Rod^Δ/Δ^ cells with active CenpE. The initially equatorial centromeres maintain their equatorial position like in RPE1 cells (Fig. 2B and C). Unexpectedly, the number of monooriented chromosomes does not increase during later prometaphase in Rod^Δ/Δ^ cells with inhibited CenpE (Fig. 2C, Rod^Δ/Δ^ CenpE-inh). Thus, unlike in RPE1 cells, CenpE appears to be dispensable for maintaining equatorial positions of already congressed centromeres. A possible explanation is that the equatorial position is normally maintained via a tug-of-war between CenpE and dynein within the kinetochore corona. As dynein is not recruited to the kinetochores in Rod^Δ/Δ^ cells, CenpE is no longer needed to counteract the dynein-produced poleward force.

### Kinetochore corona facilitates rapid relocation of the peripheral centromeres towards the spindle center

Investigations into chromosome congression usually focus on the distribution of chromosomes along the spindle axis. However, we find that during early prometaphase a larger-scale changes occur in the orthogonal direction. Shortly after NEB, centromeres gather around the spindle axis within a circular area with ~4 µm radius. This compaction of the centromere distribution arises primarily from relocation of the “peripheral” centromeres that are initially positioned > 4.5 µm from the axis (Fig. 3A). The mean distance to the spindle axis for this population decreases from 6.0 ± 1.1 μm at NEB to 3.5 ± 0.8 μm in the first 300 seconds (Fig. 3B, RPE1) while the “inner” centromeres, initially residing < 4.5 µm from the axis, change their positions only slightly (Fig. 3B, mean distance 3.0 ± 1.0 μm at NEB vs. 2.9 ± 1.0 μm at 300 s). Inhibition of CenpE does not affect the dynamics of centromere gathering (Fig. 3B, RPE1-CenpE-inh). In contrast, the inward relocation of peripheral centromeres is delayed in Rod^Δ/Δ^ cells irrespective of CenpE activity (Fig. 3B). In Rod^Δ/Δ^, the mean distance to the spindle axis remains relatively constant for ~100 seconds and then decreases slower than in RPE1 (Fig. 3B). The inner centromeres also exhibit a different behavior in Rod^Δ/Δ^ cells. The mean distance for these centromeres increases from 3.0 ± 1.0 μm at NEB to 3.6 ± 1.2 μm at 90 seconds and subsequently decreases to its NEB value 300 seconds later (Fig. 3B, Rod^Δ/Δ^ and Rod^Δ/Δ^ CenpE-inh). Together, these observations suggest that the kinetochore corona but not a plus-end directed motor activity at the kinetochore brings the initially scattered centromeres closer to the spindle axis.

**Fig. 3.**
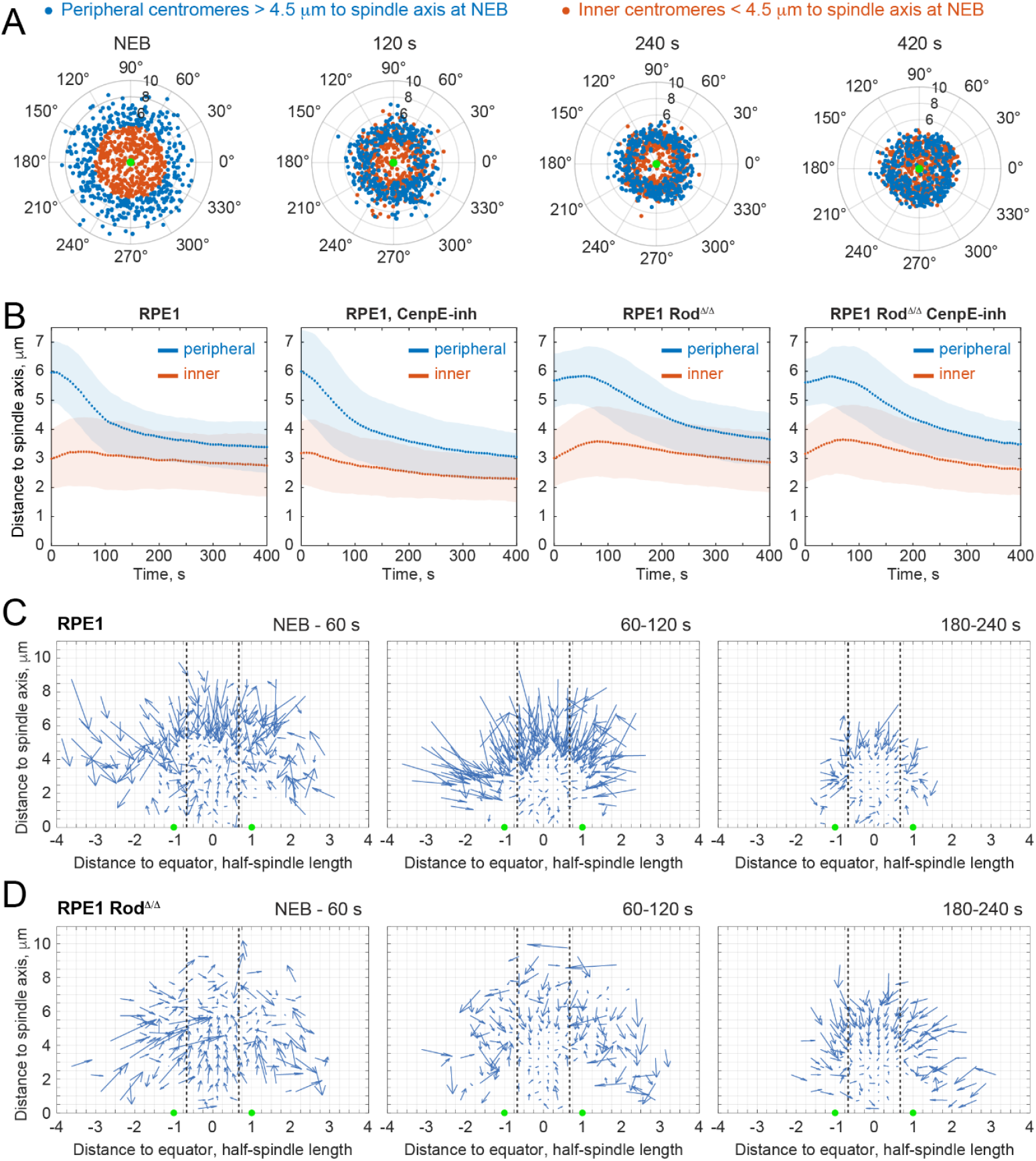
Kinetochore corona facilitates rapid convergence of the peripheral centromeres towards the spindle center. **(A)** ρ-θ distribution of centromeres during prometaphase in RPE1 cells (876 centromeres in 19 cells) at various times of prometaphase. Notice rapid (<240 s) gathering of the peripheral centromeres (blue) in a ring around the spindle axis. **(B)** Mean (with STD) distance from the spindle axis to the “peripheral” (blue) and “inner” (orange) centromeres during prometaphase. CenpE activity is inhibited with 20-nM GSK923295 (CenpE-inh). **(C)** Quiver plots showing the direction and amplitude of centromere movements at various times and within different parts of the spindle. Each arrow depicts the mean position change for all chromosomes within a small spatial bin over (marked by grey lines). Dashed black lines mark monoorientation zones of the spindle. Notice prominent near-synchronous centripetal movement of centromeres in RPE1 and lack of similar movements in Rod^Δ/Δ^ cells.

To characterize how the scattered chromosomes gather closer to the spindle axis we split the Z-ρ spindle map of centromere positions into small spatial bins and calculate the mean displacement vectors for all centromeres within each bin at various times of prometaphase. This approach reveals a striking difference in the predominant direction of centromere movements in RPE1 vs. Rod^Δ/Δ^ cells (Fig. 3C and D). In RPE1 cells, peripheral centromeres synchronously move towards to the center of the spindle during early prometaphase (1-2 minutes after NEB, Fig. 3C). In contrast, centromeres in Rod^Δ/Δ^ cells display no predominant direction during this stage of spindle assembly. At later times (> 3 min after NEB), centromere movements become more organized, but their magnitude is consistently lower than in RPE1 cells (Fig. 3C). Thus, activities associated with the kinetochore corona determine directionality, timing, and the magnitude of centromere movements during early prometaphase.

### Centromere velocity varies with the distance to the spindle center, and it depends on the presence of the kinetochore corona

To gain insight into the nature of forces that position chromosomes prometaphase we compare velocities of centromere movements at various times and in various parts of the spindle. In RPE1 cells, the mean speed of centromeres peaks at 2.0 ± 1.4 μm/min 60-90 s after NEB and subsequently decreases to 1.5 ± 1.0 μm/min (Fig. 4A). In Rod^Δ/Δ^ cells, the mean speed remains constant throughout prometaphase at 1.4 ± 0.8 μm/min (Fig. 4A). These differences indicate that the lack of the kinetochore corona primarily affects movements of centromeres during the early stages of spindle assembly, prior to the formation of amphitelic attachments. To test this possibility, we split centromere trajectories to two periods: ‘pre-amphitelic’ and ‘amphitelic’. The time of amphitelic attachment formation is detected with the previously established criteria (9). We find that most of pre-amphitelic movements are slow at a 1.2 μm/min median speed; however, rapid movements up to 19.7 μm/min are also observed in RPE1 (Fig. 4B). Rod^Δ/Δ^ cells display the same median speed but lack the rapid movements (Fig. 4B, maximum observed speed = 7.4 μm/min). Thus, pre-amphitelic kinetochores that lack the corona lose the ability to move at higher speed. For the centromeres that have established amphitelic attachments the median speed is slightly lower in Rod^Δ/Δ^ cells (Fig. 4C). As expected, rapid movements (>7.5 μm/min) are not observed among the amphitelic centromeres in either cell type (Fig. 4C).

**Fig. 4.**
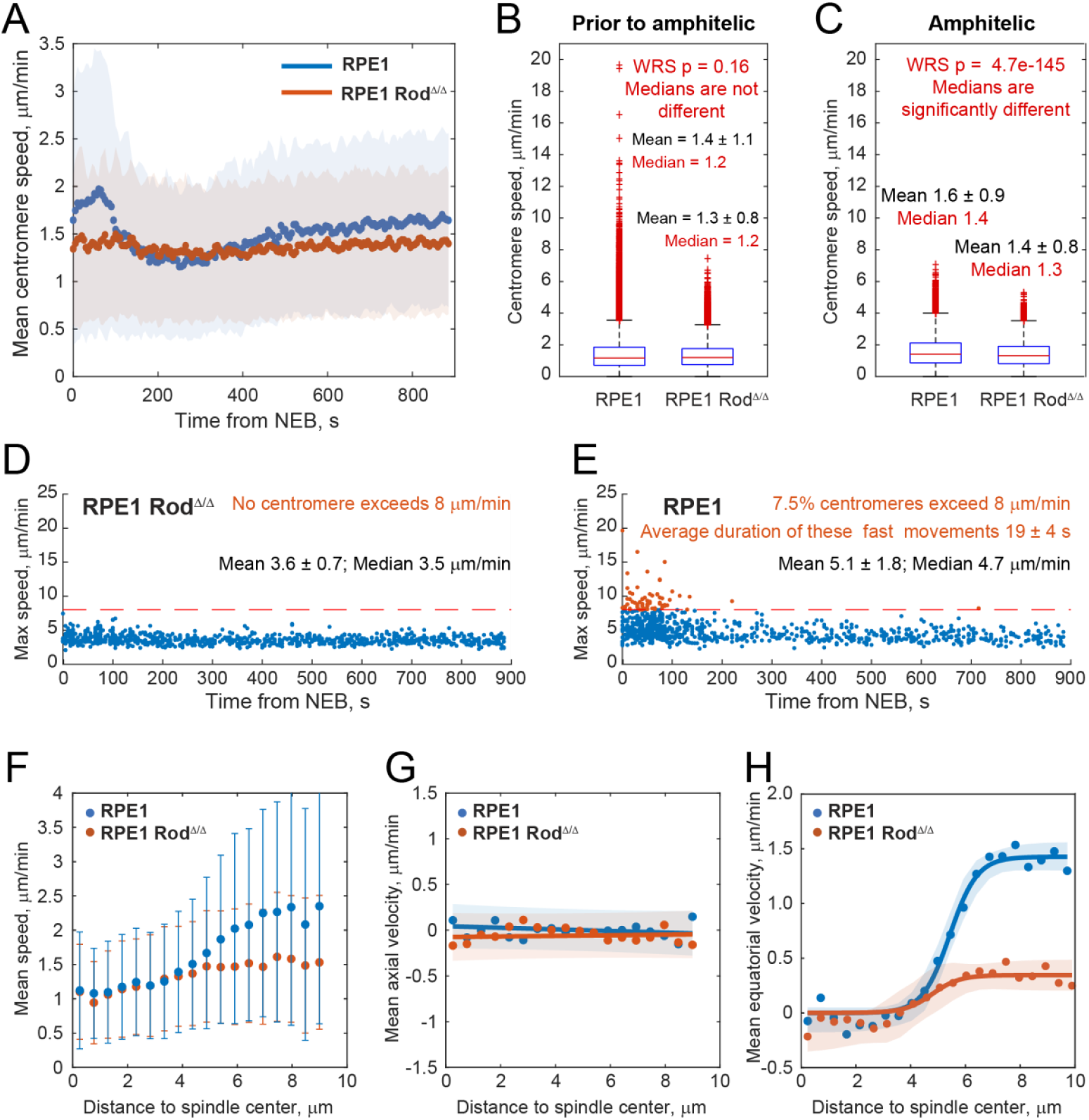
Centromere velocities in RPE1 vs. Rod^Δ/Δ^ cells at various stages of spindle assembly and in various parts of the cell. **(A)** Average speed of centromere movements (with STD) during prometaphase in RPE1 vs. RPE1 Rod^Δ/Δ^ cells. Notice that in RPE1 cells, mean centromere speed is greater in the first 60-90 s after NEB, whereas in Rod^Δ/Δ^ cells the mean velocity is constant throughout prometaphase. **(B-C)** Distribution of centromere instantaneous speed (position change over 15 s period) prior to-and after formation of amphitelic attachment in RPE1 (B) and Rod^Δ/Δ^ (D) cells. Median values are compared in the Wilcoxon Rank Sum (WRS) test. **(D-E)** Maximum speed achieved by each centromere during prometaphase in Rod^Δ/Δ^ (D) and RPE1 (E) cells. Notice that extremely rapid movements (>8 μm/min) occur infrequently during early prometaphase in RPE1 cells and not present in Rod^Δ/Δ^. **(F)** Mean centromere velocity as a function of distance to the spindle center in RPE1 vs. RPE1 Rod^Δ/Δ^ cells. Error bars are STD. Notice that in RPE1 cells, centromeres move significantly faster in the periphery (>4 μm from the spindle center). **(G-H)** Axial (G) and equatorial (H) components of centromere velocity as a function of distance to the spindle center. For the axial component (Z displacement over time), movement towards one pole is assigned a positive value and towards the opposite pole - a negative. For the equatorial component (ρ displacement over time) movements towards the spindle axis is viewed as positive and away from the axis as negative. Lines represent linear (G) or logistic (H) functions fit to the data. Shaded corridors are 95% confidence intervals for the fits.

Cytoplasmic dynein at the kinetochore corona has been implicated in the extremely rapid (up to 50 μm/min) movement of centromeres (19, 20). These movements, triggered by the initial contact between a kinetochore that lacks end-on attached MTs and an astral MT (5, 19–21) are not expected to occur in Rod^Δ/Δ^ cells. Consistent with this notion, we find that the maximum speed achieved by a typical centromere in Rod^Δ/Δ^ is less than 4 μm/min and this maximum is equally likely to be achieved at any stage of prometaphase (Fig. 4D). In contrast, centromeres achieve their maximum speed during early of prometaphase in RPE1 (Fig. 4E). At this stage, ~7.5% centromeres briefly (<15 s) exceed 8 μm/min velocity; however, most centromeres never move faster than 5 μm/min (Fig. 4E). Thus, dynein-mediated gliding of unattached kinetochores alongside astral MTs occurs infrequently.

The wide range of speeds displayed by pre-amphitelic centromeres in RPE1 prompted us to compare movements of centromeres in various parts of the spindle. In RPE1 cells, the mean speed of centromeres at the periphery (>4 μm from the spindle center) is twofold greater than in near the spindle center. In contrast, centromere mean speed is similar throughout the spindle in Rod^Δ/Δ^ cells (Fig. 4F). We also separately assess the axial (Z displacement over time) and equatorial (ρ displacement over time) components of centromere movements. For the axial component, direction toward one pole is considered as positive and toward the opposite pole as negative. We find that the mean axial velocity is near zero at all distances from the spindle center in both RPE1 and Rod^Δ/Δ^ cells (Fig. 4G), which suggests that centromeres display balanced fluctuations along the spindle axis with no preference for a particular pole. In contrast, mean equatorial velocity (positive towards the spindle axis and negative away from the axis) is nearly zero near the spindle center (<4 μm), it progressively increases at larger distances (4-7 μm), and is constant at distances >7 μm from the spindle center (Fig. 4H). In Rod^Δ/Δ^ cells, the sigmoid dependency of equatorial velocity on the distance to the spindle center is qualitatively similar but significantly less pronounced (Fig. 4H). These observations suggest that the magnitude of the force moving centromeres towards the spindle axis changes in a non-linear fashion, and this force is weaker in Rod^Δ/Δ^ cells.

The observed dependency of the equatorial velocity of centromeres may arise from the increased resistance due to gradual crowding within the inner parts of the spindle. We test this possibility with two approaches. First, we compare velocity distributions between two populations of centromeres: those residing closer to the spindle center vs. those that are >5 μm away from the center at NEB. By virtue of their initial positioning, the former move within the central part of the spindle earlier, while the latter reach the inner parts of the spindle later when the crowding is already significant. We find that both groups exhibit similar velocities at 3-5 μm distances from the spindle center (SI Appendix, Fig. S3A). In contrast, within the group of initially peripheral centromeres velocities differ significantly at larger and smaller distances to the spindle center (SI Appendix, Fig. S3B). This analysis supports the notion that the velocity is distance-but not time-dependent, i.e., the slower movement near the spindle center is not due to the gradual crowding of the space by the arriving chromosomes. Second, we test whether the dependency of the equatorial velocity on the distance is affected by inactivation of the ‘spindle ejection force’ mediated by chromokinesins KID (kinesin-10) and Kif4A (kinesin-4) (22). Simultaneous depletion of both motors via siRNA in RPE1 prevents the exclusion of chromosomes from the central part of the spindle during prometaphase (SI Appendix, Fig. S3C). Further, chromosome arms that normally orient in a radial array with their telomeres pointing outward from the spindle axis, become disorganized (SI Appendix, Fig. S3D). This change in the spindle architecture increases crowding of the inner spindle regions. Yet, no significant changes are observed in the mean centromere speed or the axial and equatorial velocities at various distances from the spindle center (SI Appendix, Fig. S4E-G).

### MTs appear at kinetochores concurrently with the initiation of nuclear envelope breakdown

In mammalian cells, forces for centromere movement have been shown two apply at two locales: at the kinetochore and at the distal ends of MTs attached to the kinetochore (7, 23). Considering recent observations that MT nucleation at the kinetochore corona is required for efficient chromosome congression (15), it is possible that the latter mechanism contributes significantly to spindle assembly. This possibility is supported by the demonstration that kinetochore attachment to short non-centrosomal MTs is common during prometaphase in RPE1 cells (8). Whether kinetochores attach to short MTs early enough to play a role in the initial congression is not known. We assess when kinetochores begin to interact with MTs by correlative light/electron microscopy (CLEM) of cells during the process of nuclear envelope breakdown.

Live cell recording of RPE1 cells (55 recordings) with labelled chromosomes and nuclear envelope demonstrate that disassembly of the nuclear envelope consistently initiates near the equatorial plane of the nucleus. Multiple small fenestrae appear in a seemingly random pattern around the nuclear perimeter. The number and size of these fenestrae increase and ~3 minutes later only remnants of the nuclear envelope are seen around the central part of the forming spindle (Fig. 5A). Concurrently with fenestration of nuclear envelope, chromosomes adjacent to the fenestra initiate directional movements. At this time, kinetochore positioned deeper inside the nucleus are still shielded by the remnants of the nuclear envelope (Fig. 5A).

**Fig. 5.**
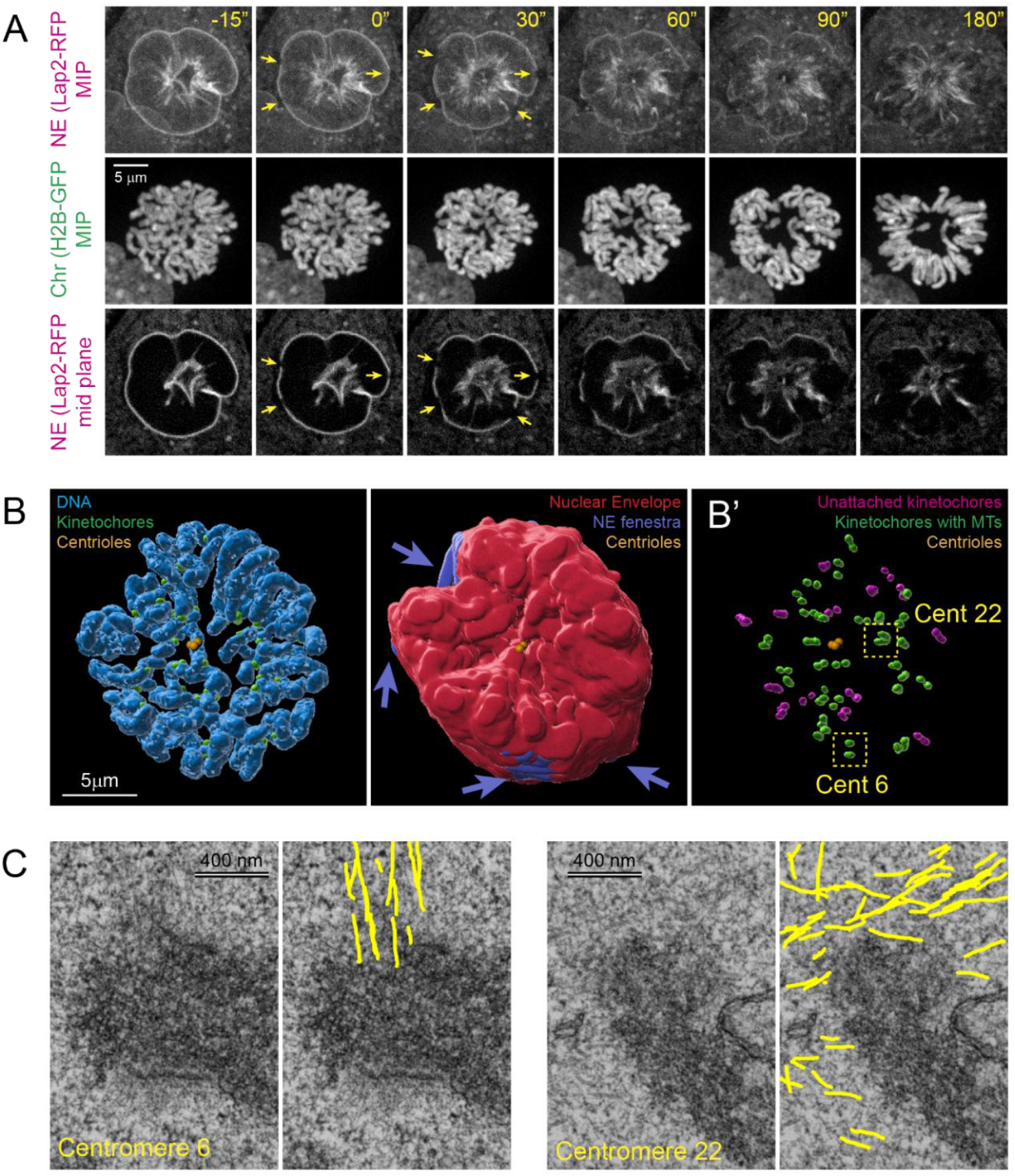
Non-centrosomal microtubules appear at kinetochores concurrently with the breakdown of nuclear envelope. **(A)** Nuclear envelope breakdown in RPE1 cells. Nuclear envelope (NE) is labeled via Lap2b-RFP and chromosomes (Chr) – via H2B-GFP. Maximum-intensity projections (MIP) of the entire cell are shown in the upper two rows. The bottom row depicts a single focal plane in the middle of the cell. Arrows point at the fenestrae that initiate NE breakdown. Time is in seconds from the formation of the first fenestra. **(B)** Correlative LM/EM reconstruction of chromosomes (blue), centrioles (orange), kinetochores (green), and NE (red) in an RPE1 fixed during NEB. Arrows point at fenestrae in the NE. Notice that all fenestrae are distant and not in direct line of sight from the spindle poles (centrioles). **(B’)** Positions of centromeres with (green) or without (magenta) MTs in the immediate proximity of at least one of the sister kinetochores. **(C)** Examples of microtubule attachment to kinetochores positioned near a fenestra (Centromere 6, boxed in B’) or deep inside the nucleus (Centromere 22, boxed in B’). Notice both end-on and lateral interactions between the kinetochores and MTs.

Guided by the time course of NEB visualized in the live-cell recordings, we fixed six RPE1 cells with GFP-labeled kinetochores and centrioles at the earliest signs nuclear envelope breakdown. In three of these cells, the nuclear envelope was completely intact, and no MT was detected within the nuclear volume (not shown). In the other three cells, several fenestrae of various size are present in the nuclear envelope (SI Appendix, Fig. S4A). Consistent with the live-cell recordings, these fenestrae are near the equatorial plane of the forming spindle. The centrosomes reside within deep invaginations of the nuclear envelope on the ventral and dorsal sides of the nucleus (Fig. 5B, SI Appendix, Fig. S4A). Unexpectedly, we find MTs associated with some kinetochores irrespective of whether these kinetochores are near a fenestra or deep inside the nucleus (Fig. 5B and B’). Both end-on attached and laterally interacting MTs are present (Fig. 5C, SI Appendix, Fig. S4B). The number of centromeres with MTs attached to at least one of the sister kinetochores varies among the three reconstructed cells (32 of 47; 23 of 46; and 1 of 46) and this number is larger in cells with larger fenestrae. In the earliest cell that contained just two small fenestrae, the only centromere associated with MTs is adjacent to a fenestra (SI Appendix, Fig. S4C and D). Thus, non-centrosomal MTs begin to appear at kinetochores concurrently with the initiation of nuclear envelope fenestration and the number of kinetochores attached to short non-centrosomal MTs increases rapidly during NEB (prior to the initiation of centromere movements).

### Mechanistic model for the early chromosome congression via dynamic interactions between the spindle and MTs at the kinetochore

Our analyses of centromere behavior reveal two prominent features of the early-prometaphase chromosome congression: 1) Centromeres move predominantly inward towards the spindle center rather than towards a spindle pole; 2) Mean velocity of the inward centromere movement is constant far from the spindle center, decreases at intermediate distances (between 7 and 4 μm), and the inward movement ceases when centromeres are closer to the center. Further, we find that many, potentially most, kinetochores are already attached to skMTs when they encounter MTs of the spindle proper. Thus, the force that moves the centromere can act on the kinetochore or at the distal ends of skMTs. These findings prompt an exploration of various kinetochore-MT interactions that explain the observed centromere behavior.

Our observations that centromeres move roughly orthogonal to the spindle axis suggest that the movement is driven by a balanced interaction with MTs that emanate from both spindle poles. For the centromere to move towards the spindle axis (inward), these forces must be directed toward the minus ends of spindle MTs. Consistent with this expectation, parameters of the initial congression are not affected by the inhibition of CenpE, the only plus-end directed motor at the kinetochore (Fig. 2B and 3B).

To evaluate the character of MT interactions capable of gathering the scattered centromeres near the equator with the dynamics observed in RPE1 cells we construct a series of computational models (SI Appendix, Supporting text). In the first model, kinetochores form stable (infinite-duration) connections to the walls of MTs produced by the spindle poles. This model stems from the observations of dynein-mediated lateral gliding of kinetochores along astral MTs (19, 20). Simultaneous interactions with MTs from the opposite spindle poles yield constant forces directed towards the minus ends of connected MTs which effectively ‘reels in’ the attached MTs (Fig. 6A). As a result, centromeres would move towards the spindle axis (Fig. 6A’) akin to lifting an object by pulling on two ropes fixed at two points on the ceiling. An essential condition in this model is the ability of MTs to pivot at the centrosome into an antiparallel configuration (24–26). However, the inward centromere velocity is predicted to be lower at larger distances from the spindle center, as the angle between the opposed MTs changes at a lower rate closer to the axis (Fig. 6A”, SI Appendix, Supporting text). The change in the angle between the vectors of the opposing forces translates the constant reeling-in rate in the periphery of the spindle into a faster movement near the spindle axis. Thus, centromere behavior observed in cells during early prometaphase cannot be explained by stable interactions between sister kinetochores and MTs from the opposite spindle poles.

**Fig. 6.**
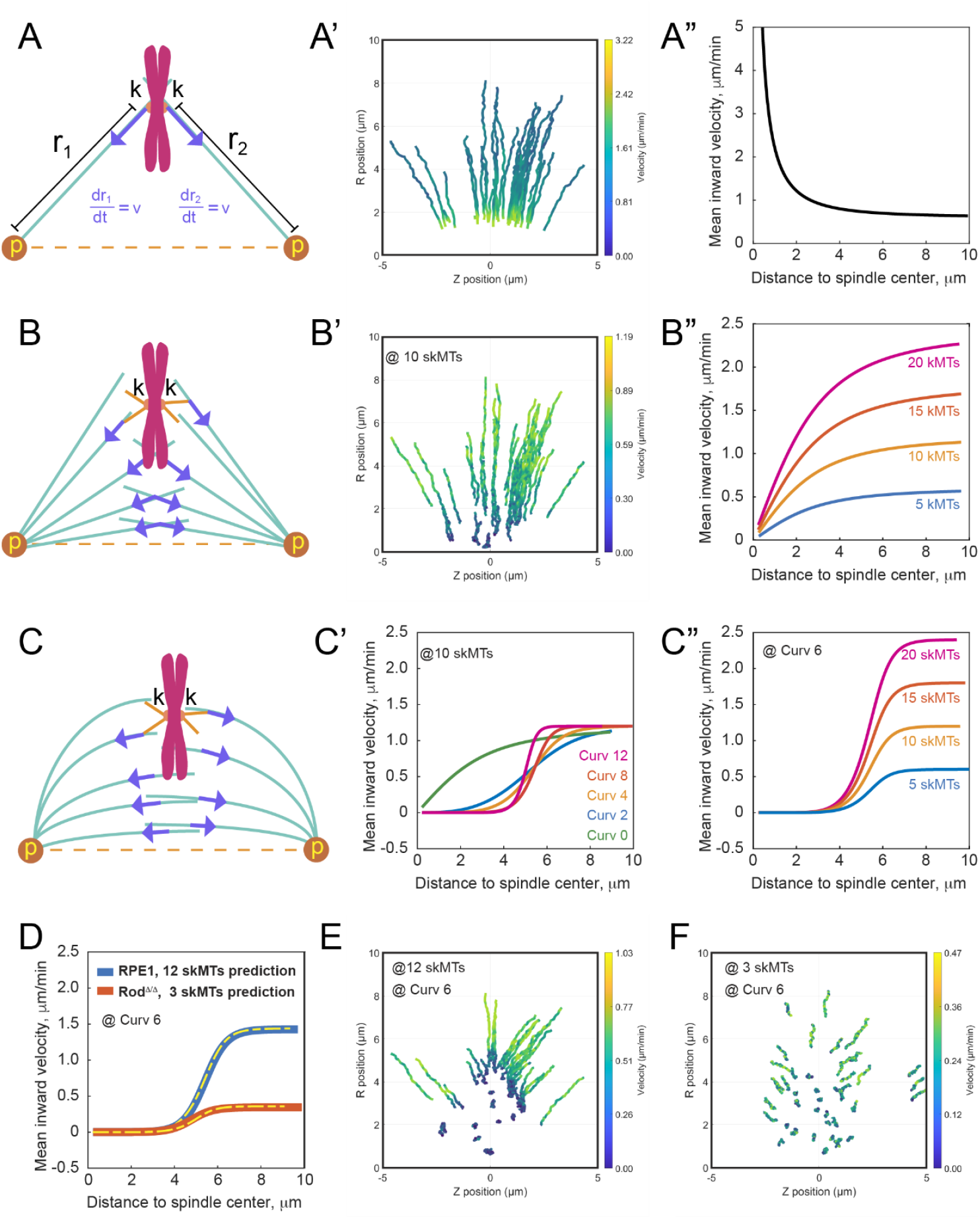
Mechanistic models for the early prometaphase chromosome congression. (A-A”) Stable interaction with spindle microtubules. (A) Kinetochores (k) form stable connections with long MTs (green lines) originating from the spindle poles (p). The force produced by the interaction with a single MT at the kinetochore plate (blue arrows) is directed towards the minus end of the MT residing at the spindle pole. Interaction with multiple MTs originating from both spindle poles move the centromere inward. (A’) Early-prometaphase (0-4 min) centromere trajectories predicted by the model. (A”) Predicted mean centromere inward velocity at various distances from the spindle center. **(B-B”) Transient interactions with straight spindle microtubules**. (B) Minus end-directed forces are generated at the distal ends of short MTs protruding from kinetochores (skMTs, orange lines). Each interaction produces a force directed towards the minus end of the spindle MT. Individual encounters between MTs are brief (seconds) but multiple MTs protrude from each kinetochore, and these MTs interact with different MTs within the spindle. (B’) Early-prometaphase centromere trajectories predicted by the model. (B”) Predicted mean centromere inward velocity at various distances from the spindle center for various skMTs numbers. (C-C”) **Transient interactions with curved spindle microtubules**. (C) Similar to (B) but MTs are curved to mimic the shape of the spindle. (C’) Effects of MT curvature on the mean centromere inward velocity at various distances from the spindle center (at constant skMT number). (C”) Effects of skMT numbers on the mean centromere inward velocity at various distances from the spindle center (at constant spindle MT curvature). **(D)** Mean centromere inward velocity at various distances from the spindle center observed (solid lines) in cells and predicted (dashed lines) by the model outlined in (C). **(E-F)** Early-prometaphase centromere trajectories predicted for kinetochores with 12 (E) or 3 (F) skMTs by the model outlined in (C). All other parameters of the model are constant.

As an alternative, we considered live-cell observations of transient dynein-mediated interactions at the distal ends of skMTs and spindle MTs (7, 8, 23). Chromosomes propelled by these interactions exhibit characteristic ‘jerks’ rather than smoother linear movements (7, 8) which indicates that the force of constant magnitude is exerted randomly on the short timescale (seconds). In this scenario, congression is driven by numerous transient connections to multiple long MTs that are within the reach of skMTs (Fig. 6B, SI Appendix, Supporting text). The transient-interaction model predicts that centromeres move predominantly inward, and their velocity is greater at larger distances from the spindle center (Fig. 6B’). Yet quantitatively the velocity-distance function predicted by the model differs significantly from the sigmoid dependency observed in cells (Fig. 6B”).

Our previous findings that the shape and architecture of the spindle determine the time and place of amphitelic attachment formation (9) inspired us to explore the transient-interaction model in the context of a more realistic geometry of spindle MTs. Chromosome congression has been shown to be less efficient in cells that lack the MT crosslinker PRC1 (9, 27). We reasoned that without the cross-links, MTs are less likely to bend towards the equator and tend to grow in more normal directions to the spindle axis. Analyses of centromere movements in PRC1-depleted cells reveal no significant changes in the total speed or mean velocity along the spindle axis (SI Appendix, Fig. S5 A and B). However, the dependency of the equatorial velocity on the distance to the center changes significantly: the inward movement is slower at intermediate distances but at larger distances from the spindle center the velocity is significantly higher than in untreated RPE1 cells (SI Appendix, Fig. S5C). These effects support the notion that the shape of spindle MTs must be considered in modeling of chromosome behavior.

The modified transient-interaction model with MTs curved along ellipsoidal surfaces that resemble the shape of prometaphase spindle (Fig. 6C, SI Appendix, Supporting text) correctly predicts a sigmoid dependency of the inward velocity on the distance to the spindle center. The exact shape and magnitude of the predicted function depend on multiple parameters (SI Appendix, Model Details and Derivations) with the curvature of spindle MTs (Fig. 6C’) and the number of short MTs protruding from the kinetochores (Fig. 6C”) being particularly important. Guided by our recent description of the spindle shape (9) and the numbers of MTs observed by EM at RPE1 kinetochores prior to amphitelic attachment formation (8) we find a combination of these parameters that predict the velocity-distance function that matches the one observed in RPE1 cells (Fig. 6D). At these parameters, the model predicts that the centromeres predominantly move towards the spindle center, but their inward movement ceases when they reach within 3-4 μm from the spindle axis (Fig. 6E). Further, the model predicts a steady decrease in the number of polar (inherently monooriented) chromosomes at 2.7 ± 0.9 per minute rate, which accurately matches our experimental observations (Fig. 2B, RPE1). Thus, the dynamics of chromosome congression in RPE1 cells can be explained by transient (seconds) interactions between ~12 skMTs and antipolar arrays of properly shaped spindle MTs.

We then explore whether our computational model can explain the changes in the pattern and velocity of centromere movements observed in Rod^Δ/Δ^ cells. Astoundingly, we find that decreasing the number of skMTs from 12 to 3 with no change in other parameters yields a predicted centromere velocity-distance function that perfectly matches the experimental function observed in Rod^Δ/Δ^ cells (Fig. 6D). Further, predicted trajectories (Fig. 6F) are consistent with the lack of directionality in centromere movements observed in Rod^Δ/Δ^ (Fig. 3D). Furthermore, the rate of monoorientation correction is predicted to be 1.5 ± 0.6 chr/min, which matches the experimentally observed (Fig. 2B, Rod^Δ/Δ^). Thus, the numerous abnormalities observed in cells lacking the RZZ complex can be explained solely by the lower number of skMTs due to the lower efficiency of MT nucleation at the kinetochores (15, 17).

## Discussion

Inefficient chromosome congression, manifested by the increased number and persistence of monooriented chromosomes, is perhaps the most common abnormality of cell division (11). Numerous molecular deficiencies have been linked to hindered congression; however, mechanistic understanding of how the initially scattered chromosomes rapidly converge in a tight plate near the equator is lacking (11, 28). One impediment to understanding congression mechanisms is that the typical behavior of chromosomes during early stages of spindle assembly has not been quantitatively characterized. Here we use an unbiased population-level analysis of centromere movements in various parts of the spindle. Three characteristic features of early-prometaphase chromosome behavior emerge from these analyses: 1) Centromeres initiate directional movements concurrently with the nuclear envelope breakdown; 2) Scattered centromeres predominantly move directly towards the center of the spindle and rarely towards a spindle pole. Indeed, the number of monooriented chromosomes is maximal at NEB; 3) Velocity of the centripetal centromere movement decreases as the centromere approaches the spindle axis. These features are not consistent with the chromosome behavior envisioned in the current models of spindle assembly that stem from the ‘Search and Capture’ hypothesis (29).

It is commonly assumed that prior to their attachments to MT plus ends, kinetochores glide alongside MTs via forces generated by MT motors within the kinetochore’s outer plate. CenpE-dependent movements along MT bundles towards the plus ends as well as remarkably rapid dynein-dependent minus-end directed gliding along single MTs have been directly observed in cells (19, 20, 28, 30). Consistent with the latter observations, we also detect rapid (>8 μm/min) kinetochore movements in the wild-type RPE1 but not in Rod^Δ/Δ^ cells that lack dynein at their kinetochores (Fig. 4 B and C). However, these rapid dynein-dependent movements are infrequent (7-8% of chromosomes) and thus they are unlikely to make a major contribution to the initial congression.

Strong evidence exists that due to differences in post-translational modifications of tubulin within astral MT vs. less dynamic MT bundles, dynein engages upon the initial interaction with an astral MT (31, 32). Thus, kinetochores are expected to move initially towards a spindle pole. Monoorientation, created by this early dynein activity, is subsequently corrected by the CenpE-mediated movement to the plus ends of MTs near the spindle equator (32). In this scenario, lack of dynein at the kinetochore should suppress monoorientation, while lack of CenpE activity should impede congression of monooriented chromosomes (11, 33). Yet, our systematic analyses of centromere movement during early prometaphase demonstrate that inhibition of CenpE does not impede relocation of centromeres closer to the center of the spindle (Fig. 2 and 3, CenpE-inh). In contrast, chromosome movements towards the spindle center are impeded in cells that lack dynein at the kinetochores (Fig. 2 and 3, Rod^Δ/Δ^). These effects are the opposite of the expected for centromere movements driven by motors residing within the kinetochore and asynchronous capture of astral MT by sister kinetochores. Thus, while some chromosomes clearly follow the “classic” monoorientation-congression pattern during prometaphase, this behavior is not common.

Two intriguing features of centromere movements during early prometaphase are the predominant centripetal direction of the movement and the sigmoid dependency of centromere velocity on the distance from the center. We test several models that could explain these features and arrive at a single plausible mechanism. Common in all tested models is that a centromere with two sister kinetochores engages MTs from both spindle poles. This feature is necessary to explain the generally centripetal direction of the movement. However, a nearly synchronous capture of MTs from both spindle poles is possible only for unrealistically large kinetochores (34, 35). For example, if ~1,000 MTs splay out from each centrosome and the nuclear volume is roughly a cylinder with 4 μm height 10 μm radius, then, at the cylindrical surface farthest away from the spindle center, a single MT contact requires no less than 0.5 x 0.5 μm^2^ area, which is significantly larger than the plate of a human kinetochore (36). The insufficient size problem would be resolved, if spindle MTs interact not with the kinetochore plate directly but with multiple short (<1μm) pivoting MTs protruding from the plate.

Our live-cell and CLEM data suggest that laterally as well as end-on attached MTs begin to accumulate at kinetochores concurrently with the breakdown of nuclear envelope, prior to the initial contacts with spindle MTs (Fig. 5). These data are consistent with previous reports that the majority of RPE1 kinetochores (>75%) are in close contact with ~30 non-centrosomal MTs, >10 of which appearing to attach end-on during mid-prometaphase (8). Thus, a typical kinetochore appears to bear multiple skMTs during its initial encounter with spindle MTs. A corollary of this feature is that spindle microtubules arriving near a centromere are more likely to interact with the protruding skMTs rather than with the kinetochore plate.

Distal ends of skMTs have been observed to rapidly connect to the walls of adjacent microtubules within the spindle and be pulled toward the minus ends of the spindle microtubules (7, 23) in a characteristic ‘jittery’ pattern in which low-amplitude momentary poleward jumps are intermittent with short pauses (7). In contrast, kinetochore gliding alongside of astral MTs driven by dynein within the kinetochore outer layer are smooth (5, 21, 31). These observations support the idea that while the skMTs-MTs interaction produces a constant-magnitude net force, this force arises from the rapid attachment-detachment cycle exhibited by individual skMTs. Our computational analyses demonstrate that this type of interaction correctly explains centromere movements observed during early prometaphase (Fig. 6). Continuous gliding of kinetochores on intersections of astral MTs results a wrong spatial distribution of centromere velocities and slower decrease in the number of monooriented chromosomes (Fig. 6A’ and A”).

Formally, a gradual slowing down of centromeres’ inward movements could be caused by a distance-dependent ejection force acting on the chromosome arms (37, 38) even in the case of stable interactions between kinetochores and astral MTs. However, we find that the velocity-distance function does not change in cells depleted for both kinesin motors responsible for the ejection of chromosome arms (SI Appendix, Fig. S3F). Thus, the ejection force is not a major factor in the slower movement of centromeres near the spindle center. Neither our analyses support the possibility that the slower movements in the inner parts of the spindle are due to crowding of the space by the arriving chromosomes (SI Appendix, Fig. S3A and B). Ruling out these formal possibilities, leaves the transient nature of interactions with spindle MTs as the only viable reason for the observed velocity-distance relationship.

While the prediction of the transient-interaction model that centromeres move slower near the spindle center holds for a wide range of MT distributions, matching the observed sigmoid velocity-distance dependency that MTs emanating from the spindle poles are curved along ellipsoidal surfaces that resemble the natural shape of the spindle (Fig. 6C and C”). Remarkably, this modification of the model yields predicted distance-velocity functions that are indiscernible from the experimentally observed (Fig. 6D). Further, the prominent difference in the distance-velocity functions in RPE1 vs. Rod^Δ/Δ^ arises from changing a single parameter of the model – the number of short MTs protruding from the kinetochore. Importantly, a lower number of short MTs at the kinetochore is expected as lack of dynein and more specifically its light intermediate chain at the kinetochore is known to suppress nucleation of MTs at the kinetochore corona (15).

Based on these observations, we propose that early-prometaphase behavior of chromosomes in human cells is principally determined by transient interactions between numerous short MTs protruding from the kinetochores and properly shaped and distributed spindle MTs (Fig. 7). These interactions, mediated by multivalent dynein complexes, result in rapid, short, and random, excursions of skMTs towards the minus ends of spindle MTs that radiate from both poles. Transient relatively weak interactions have been shown to be surprisingly impactful drivers of motion at the molecular scale in other contexts (39). Noteworthy is also that a single dynein motor is capable of generating forces that are an order of magnitude greater than the minimal force required for moving a chromosome (~0.1 pN, see reference (40)). An important advantage of brief contact in a repeating detachment-reattachment cycles is that centromeres move toward the spindle center by making numerous contacts with various spindle MTs, rather than pulling along the same MTs all the time. Consequently, the direction and magnitude of the resultant force changes in different parts of the spindle. One potentially important function of the resulting telescopic velocity is synchronized arrival of centromeres to the area with high concentration of antiparallel bundles facilitating the efficient formation of amphitelic attachments (9).

**Figure 7.**
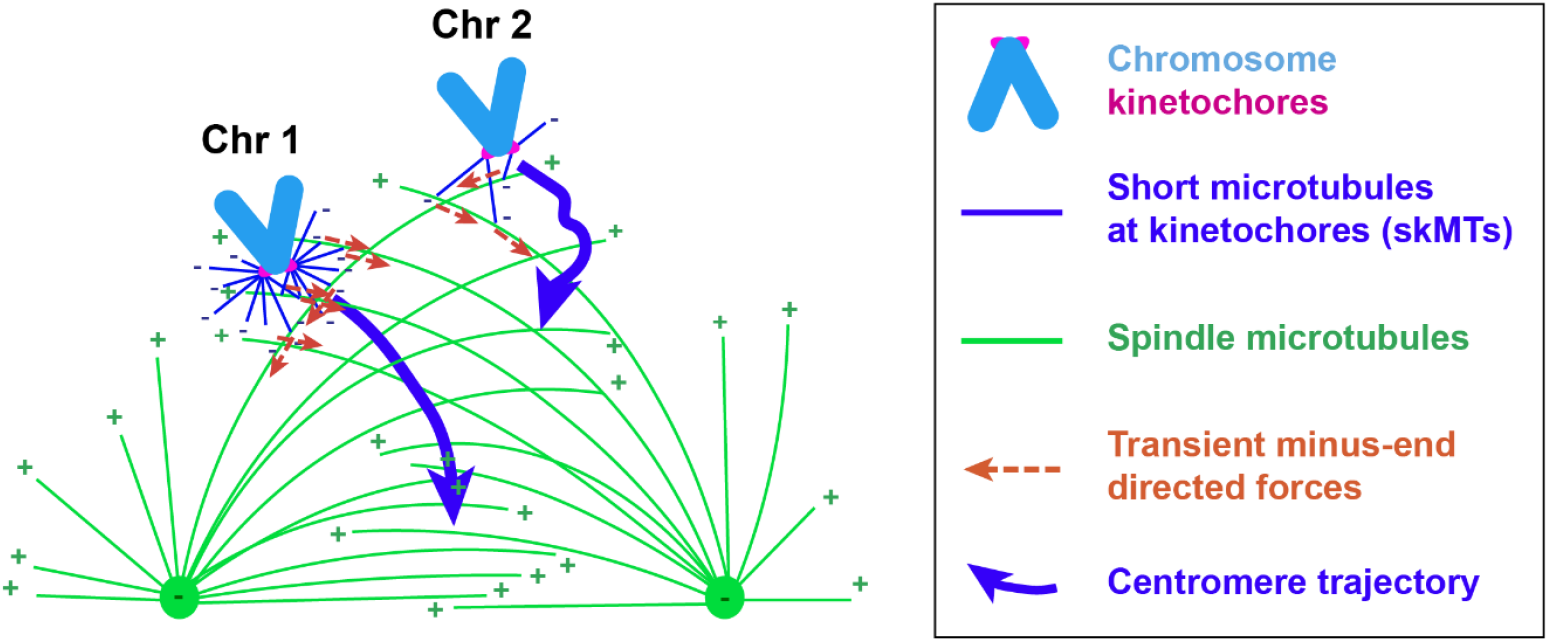
Mechanism for centromere centripetal movements during early stages of spindle assembly. Gathering of the initially scattered chromosomes onto the equatorial plate during is driven primarily by transient interactions between skMTs protruding from kinetochores and curved MTs of the spindle (“Chr 1”). These interactions involve dynein that acts at the minus ends of skMTs. Magnitude of the resultant force and direction of centromere movement depends on the number of interactions (in turn determined by the number of skMTs) and the shape of spindle MTs. A sufficient number of skMTs in the context of properly shaped spindle MTs moves the centromere towards the spindle center (Chr1). The velocity deceases as the angle between the centromere trajectory and spindle MT changes. An insufficient number of skMTs, e.g., in cells with suppressed MT nucleation at the kinetochore (Rod^Δ/Δ^) results in slower movement and loss of directionality (“Chr 2”).

The model we propose places directionality and density requirements on the MT network of the spindle. It is now increasingly apparent that distinct populations of MTs in the spindle exist (41), and whether this complex network of MTs (1) and KTs (42) in the prometaphase spindle corresponds to these requirements remains to be seen. Other interesting questions are how MT branching (43) viscoelastic nonlinear mechanics of MT network (44) and chromosome arms (45) could affect centromere convergence. Also, in other contexts, MT configurations can be optimized for cargo capture (46), and it would be interesting to investigate the optimal configurations of MTs for rapid chromosome arrival to the spindle. Importantly, the model produces accurate numeric predictions not just the distance-velocity function but also for the rates of monoorientation correction.

Formally, in the presence of skMTs, interactions among these MTs can result in chromosomes converging on their center-of-mass independent of the forces on spindle MTs. Subsequently, the spindle poles could be pulled into positions defined by the chromosomes (47). However, computational analysis of the relationships between the positions of spindle poles and orientation of the forming chromosome plate demonstrates that movements of the spindle poles during early prometaphase are not responsive to changes in the positions of centromeres. Therefore, the position and orientation of the spindle during prometaphase are not determined by the distribution of chromosomes (SI Appendix, Supporting text). Instead, the centromeres’ center-of-mass seems to follow the movements of the spindle center whose position in turn is determined by the positions of the centrosomes. Further, the model that considers direct interactions among the centromeres predicts complex, non-monotonic, and unphysical velocity/distance dependence. Thus, we rule out the formal possibility of chromosome congression via centromere-centromere interactions.

Another possibility that must formally be considered is that chromosomes initially gather closer to the spindle center via MT-independent mechanisms. Several studies suggest than contraction of actomyosin networks (48, 49), like a fishnet, can bring chromosomes closer to the spindle. This possibility is especially relevant because the telescopic inward velocity emerges naturally in the actomyosin networks (50). However, the idea of a MT-independent mechanism is not consistent with the dramatic changes in centromere velocities and the rate of monoorientation correction observed in Rod^Δ/Δ^ cells. While these observations do not rule out involvement of actomyosin contractility in the initial congression, this mechanism is likely to be a relatively minor part of the early-prometaphase chromosome congression at most. Thus, the model described in Fig. 7 appears to offer the most reasonable mechanism of congression in human cells.

## Acknowledgments

This work was supported by the National Institutes of Health NIGMS grant GM130298 (A.K.), National Science Foundation grants DMS 1953430 (A.M.) and CAREER DMS-2339241 (C.M.) and NSF/NIGMS grant DMS-2451263 (C.M.)

## Materials and Methods

### Cell lines and chemical

RPE1 cells co-expressing Centrin1-GFP and CenpA-GFP (12), RPE1 Rod^Δ/Δ^ co-expressing Centrin1-GFP and CenpA-GFP (9), RPE1 co-expressing H2B-Neon and mRFP-LAP2b (a kind gift from MD. David Pellman, Dana-Farber Cancer Institute), RPE1 expressing Sh-PRC1 and RPE1 co-expressing Centrin1-YFP and CenpA-GFP (generated in this study) were cultured in antibiotic-free DMEM/F-12 (Gibco) medium supplemented with 10% fetal bovine serum (FBS, Gibco) at 37°C, 5% CO_2_. Culture media for RPE1 Rod^Δ/Δ^ cells were additionally supplemented with 1-mM sodium pyruvate (Gibco). CenpE was inhibited by 20-nM GSK-923295 (MedChemExpress), added to the growth medium 0.5-2.5 h prior to initiation of live cell recordings.

### Transfection

To generate RPE1 cell line co-expressing Centrin1-YFP and CenpA-GFP, RPE1 cells (Clontech) were transfected with lentivirus construct as previously described (12). Briefly, cells were first transfected with CenpA-GFP in LentiLox 3.1. Lentivirus were added to the standard culture media with 1:100 Polybrene (TR-1003-G; EMD Millipore) for ~12 hr. Individual clones selected for the high expression level of GFP labelling both centromeres and chromosome arms were subsequently transfected with centrin1-YFP. Stable lines were then selected by microscopy screening of individual clones generated by limited dilutions.

siRNA transfection was performed with the following target sequences: 5’-AAGCGCGCTTTGTAGGATTCG-3’ for Kid (38) and 5’-CAGGTCCAGACTACTACTC-3’ for

Kif4a (51). siRNA oligonucleotides were co-transfected into RPE1 cells by electroporation with a Nucleofector (X-001 program; Amaxa Biosystems). Live imaging or fixation was performed 48 hr after depletion. Depletion efficiency was monitored by phenotypic analysis. Only cells that failed to form a chromosome ring during prometaphase were analyzed.

### Live-cell microscopy

Cells were grown on #1.5 glass coverslips in Petri dishes for 48-72h. One day prior to the recording, culture media was replaced with phenol-red free mixture of DMEM/F-12 containing 10% FBS and penicillin/streptomycin (P4333; Sigma-Aldrich). Approximately 3 h prior to the recordings, coverslips were mounted on Rose chambers and placed on the microscope stage. The chambers were maintained at 37.0 ± 0.3°C within a custom-built enclosure. Imaging was performed with a spinning-disc confocal scanner (Yokogawa, X1) attached to a Nikon Ti2E microscope equipped with a *λ*PlanApo 100×1.45 NA oil-immersion objective. 488-nm excitation light intensity was kept at ~10 nW/mm^2^ (~40 μW out of the lens). For tracking centromere movements, higher-temporal resolution recordings were collected every 5 seconds at 500-750 nm z steps (17-20 planes) and at 110-nm XY pixel size. With the exception of GSK-treated RPE1 Rod^D/D^ recordings, datasets from our previous paper were used in this study (Renda et al., 2022). All the other time-lapse recordings were performed at 15-30 second intervals. 640-nm and 561-nm excitation light intensities were kept at ~10 nW/mm^2^. Images were processed using SoftWoRx (GE Healthcare, USA) and IMARIS (Oxford Instruments) 3D imaging software.

### Immunofluorescence

Cells were pre-extracted in warm PEM buffer (100-mM PIPES [pH 7], 1-mM EDTA, 1-mM MgCl_2_) supplemented with 0.5% Triton X-100 for 30 seconds. Specimens were then fixed for 10 minutes in warm PEM buffer containing 3.2% paraformaldehyde (EM grade; EMS) and 0.1% glutaraldehyde (G5882; Sigma-Aldrich), followed by brief washing in PBS and reduction in an aqueous solution of 30-mM sodium borohydride (452882; Sigma-Aldrich) for 5 minutes. After reduction, cells were washed with PBS and then incubated with blocking/permeabilization buffer (PBS with 3% BSA and 0.5% Triton X-100) for 30 minutes. Microtubules were visualized with DM1a monoclonal anti-a-tubulin antibody (T9026; Sigma-Aldrich) at 1:200 followed by a secondary antibody conjugated with Alexa Fluor 647 at 1:100 (A21236; Invitrogen Molecular Probes). Both primary and secondary antibodies were dissolved in blocking/permeabilization buffer. Chromosomes were stained with Hoechst 33343 (Molecular Probes) at 1 mg/ml. Kid- and Kif4A-codepleted RPE-1 cells were lysed in warm PEM buffer (100 mM Pipes [pH 6.9], 2.5-mM EGTA, and 5-mM MgCl_2_) supplemented with 1% Triton X-100 for 1 minute before fixation with 1% glutaraldehyde (G5882; Sigma-Aldrich). Images of fixed cells were collected on the same microscope as live-cell recordings at 73 or 110-nm XY pixels and 200-nm Z-steps. All images were deconvolved with the SoftWoRx 5.0 (Applied Precision) and objective lens-specific point spread function. Precise Axial and Equatorial views of the spindle were generated by rotating the volume in 3-D to orient the spindle axis defined by the 3-D coordinates of both spindle poles. 3-D reconstructions of centromeres and chromosomes were visualized in Imaris by Surfaces and Spots material options.

### Correlative Light Electron Microscopy (CLEM)

Cells were fixed for 30 minutes in 100-mM sodium cacodylate buffer [pH 7.4] (EMS) containing 2.5% glutaraldehyde (G5882; Sigma-Aldrich). Chromosomes were stained in 100-mM sodium cacodylate buffer containing 1 mg/ml Hoechst 33342 (Molecular Probes) for 5 minutes. Complete Z-series were collected as in fixed-cell immunofluorescence preparations. EM embedding and serial sectioning were done as previously described (Rieder and Cassels 1999). 80-nm sections were imaged on a JEM 1400 microscope (JEOL) operated at 80 kV using a side-mounted 4.0-megapixel XR401 sCMOS AMT camera (AMT). Complete image series recorded at 5K magnification were used to reconstruct whole-cell volumes. These volumes were aligned with the light microscopy images by matching position of chromosome arms. Serial higher-magnification images (40K) were then collected to detail the distribution of microtubules near centromeres.

### Trajectory Processing

For each chromosome trajectory, both experimental and simulation, instantaneous velocities were calculated using finite differences **v**(*t*) = [**x**(*t* + *Δt*) − **x**(*t*)]/*Δt* over three frames, *Δt* = 15*s*. We analyzed the velocity data using two different decomposition methods. The first is in the spindle cylindrical decomposition, yielding a radial velocity (*v*_*r*_ = *dr*/*dt*) and an axial velocity (*v*_*z*_ = *dz*/*dt*). The second is a decomposition that projected the velocity vector onto two distinct axes: (1) the direction toward the spindle center, capturing convergent movement, and (2) the orthogonal component, capturing lateral movements. For this second decomposition, given a chromosome at position **x** = (*z, r*) with velocity **v**, we calculated the unit vector pointing toward the spindle center: 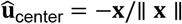. The velocity toward center was then calculated as 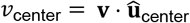, and the orthogonal velocity was calculated as 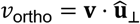. Velocities exceeding 8 *μ*m/min were identified as jumps and excluded from analysis. The filtered velocity data were binned either by radial distance from the spindle axis (*r*) or by total distance from the spindle center 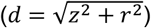 into equally-spaced bins, with mean velocities and standard deviations calculated for each bin. The spatially-dependent velocities were then fit with four-parameter logistic or two-parameter linear functions.

### Computational Models and Simulation

We implemented and simulated two distinct models representing competing hypotheses for chromosome movement in early mitosis. All simulations were performed in two-dimensional cylindrical coordinates (*z, r*) with spindle poles fixed at positions (±*L*, 0). Initial chromosome positions were sampled from experimental distributions, and simulations incorporated Brownian motion through additive Gaussian noise with a standard deviation of *σ*. A sub-timestep integration scheme (*dt* = 0.01s) was used for accurate resolution of stochastic events, with results downsampled to 5s to match experimental time resolution. The first model, the stable attachment model, assumes chromosomes maintain persistent connections to both spindle poles through microtubules. In this framework, motion is driven by the inward flux of microtubules while these attachments are maintained. The second, the transient interaction model, considers chromosome movement through repeated cycles of binding and unbinding of motor proteins to a dense network of spindle microtubules. Each chromosome was modeled with several independent force-generating elements that stochastically bind to microtubules from either pole with rates dependent on local microtubule density, and unbind with a constant probability. Within this model, we considered several regimes, including microtubule-density-limited and force generator (motor)-limited conditions. The force-generating elements in this model could follow either straight paths directly toward the poles or curved paths defined by the spindle geometry. Details are in the SI Appendix, Supporting text.

