## Supplementary Information for "The kinetochore corona orchestrates chromosome congression through transient microtubule interactions"

**This PDF file includes:**

Supporting text

Figures S1 to S5

Tables S1 to S3

### Model Details and Derivations

#### S1. Introduction and Coordinate System

This supplement provides complete mathematical details for the chromosome movement models analyzed in the main text. We work in cylindrical coordinates $(z, r)$ where $z$ represents position along the spindle axis and $r$ represents radial distance from this axis. The spindle poles are located at positions $\mathbf{p}_1 = (L, 0)$ and $\mathbf{p}_2 = (-L, 0)$, where $L$ is the half-spindle length. Throughout this document, position vectors are denoted $\mathbf{x} = (z, r)$ and velocities as $\mathbf{v} = d\mathbf{x}/dt$. Time evolution follows the general form $d\mathbf{x}/dt = \mathbf{v}(\mathbf{x}, t) + \boldsymbol{\eta}(t)$ where $\boldsymbol{\eta}$ represents stochastic noise terms.

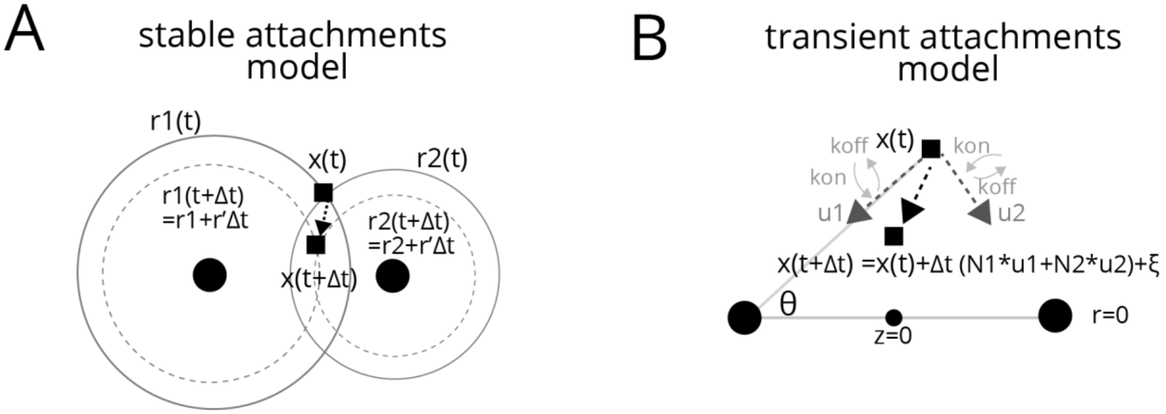


#### S2. Stable Attachment Model

##### S2.1 Biological Motivation and Assumptions

The stable attachment model represents the assumption that chromosome motion is driven by end-on attachments to microtubules emanating from opposite spindle poles. The key assumptions underlying this model are: (1) once attached, kinetochore-microtubule connections persist throughout prometaphase, (2) chromosomes remain tethered to both poles simultaneously, and (3) movement occurs through coordinated depolymerization of kinetochore microtubules, causing the effective "attachment spheres" centered at each pole to contract over time. This framework treats chromosome movement as purely geometric, determined by the intersection of two shrinking spheres.

##### S2.2 Mathematical Formulation

Consider a chromosome at position $\mathbf{x} = (z, r)$ connected to both spindle poles. The constraints imposed by stable attachments to the poles are:

$$\| \mathbf{x} - \mathbf{p}_1 \|^2 = r_1^2(t)$$
$$\| \mathbf{x} - \mathbf{p}_2 \|^2 = r_2^2(t)$$

where $r_1(t)$ and $r_2(t)$ are the time-dependent radii of the attachment spheres. Expanding these constraints:

$$r^2 + (z - L)^2 = r_1^2$$
$$r^2 + (z + L)^2 = r_2^2$$

The sphere radii evolve according to prescribed functions that can incorporate biological effects such as polar ejection forces:

$$\frac{dr_1}{dt} = v_r(r_1) + \sigma_1\sqrt{|v_r(r_1)|}\xi_1(t)$$
$$\frac{dr_2}{dt} = v_r(r_2) + \sigma_2\sqrt{|v_r(r_2)|}\xi_2(t)$$

where $v_r(r)$ represents the deterministic, distance-dependent rate of change of the sphere radius and $\xi_i(t)$ are independent white noise processes.

##### *S2.3 Velocity Derivation*

To find the chromosome velocity, we differentiate the constraint equations with respect to time:

$$\frac{d}{dt}[r^2 + (z - L)^2] = \frac{d}{dt}[r_1^2]$$
$$\frac{d}{dt}[r^2 + (z + L)^2] = \frac{d}{dt}[r_2^2]$$

Expanding the derivatives:

$$2r\frac{dr}{dt} + 2(z - L)\frac{dz}{dt} = 2r_1\frac{dr_1}{dt}$$
$$2r\frac{dr}{dt} + 2(z + L)\frac{dz}{dt} = 2r_2\frac{dr_2}{dt}$$

This linear system can be solved for the velocity components. Subtracting the first equation from the second:

$$2(z + L)\frac{dz}{dt} - 2(z - L)\frac{dz}{dt} = 2r_2\frac{dr_2}{dt} - 2r_1\frac{dr_1}{dt}$$
$$4L\frac{dz}{dt} = 2\left(r_2\frac{dr_2}{dt} - r_1\frac{dr_1}{dt}\right)$$

Therefore:

$$\frac{dz}{dt} = \frac{r_2\frac{dr_2}{dt} - r_1\frac{dr_1}{dt}}{2L}$$

Substituting back to find $dr/dt$, and after some algebra, we find

$$\frac{dr}{dt} = \frac{r_1\frac{dr_1}{dt}(z + L) - r_2\frac{dr_2}{dt}(z - L)}{2Lr}.$$

##### *S2.4 Simulation Algorithm*

The stable model simulation proceeds as follows:

1. Initialize chromosome positions from experimental data
2. Calculate initial sphere radii: $r_1 = \| \mathbf{x} - \mathbf{p}_1 \|$, $r_2 = \| \mathbf{x} - \mathbf{p}_2 \|$
3. At each timestep $dt$:
   - Update sphere radii: $r_i(t + dt) = r_i(t) + v_r(r_i)dt + \sigma_i\sqrt{|v_r(r_i)dt|}\eta_i$
   - Calculate new position using the velocity equations
   - Add Brownian noise: $\mathbf{x}(t + dt) = \mathbf{x}(t) + \mathbf{v}dt + \sigma\sqrt{dt}\boldsymbol{\eta}$

##### *S2.5 Model Variants and Parameters*

We examined two variants of the stable model:

**Constant contraction:** $v_r(r) = v_0$ , a constant depolymerization rate.

**Polar ejection forces:** $v_r(r) = v_0\big(1 - \alpha\exp(-r/r_0)\big)$, captures the repulsive forces exerted by astral microtubules near the poles (causing expansion, $v_r > 0$) which slow sphere contraction when chromosomes are far from the poles.

#### S3. Transient Interaction Model

##### *S3.1 Biological Motivation*

The transient interaction model is motivated by the hypothesis that transient kinetochore-associated or short-microtubule-associated motors can transport chromosomes along the microtubules before stable end-on attachments form. Rather than maintaining persistent connections, chromosomes in this model undergo repeated cycles of binding to and unbinding from fixed-geometry microtubules throughout the spindle.

##### *S3.2 Theoretical Foundation: Filament-Mediated Transport*

To understand how transient interactions produce directed movement, we first derive the theoretical framework for transport in a filament network. Consider spindle poles at positions $(0, \pm L)$ that emit microtubules with a specified angular distribution. Each microtubule extends from its pole, and chromosomes can bind to these filaments and be transported toward the pole.

The key insight is that the local density of microtubules determines the binding probability, while the geometry of the filament network determines the direction of transport. We define the angular distribution $f(\theta)$ where $\theta$ is measured from the pole-to-pole axis. Specifically:

- For the pole at $z = +L$: $\theta = 0$ points toward $z = 0$ (downward along the spindle axis)
- For the pole at $z = -L$: $\theta = 0$ points toward $z = 0$ (upward along the spindle axis)
- For both poles: $\theta = \pi/2$ points perpendicular to the spindle axis

##### *S3.3 Derivation of Microtubule Density*

Starting from first principles, we calculate the spatial distribution of microtubules. Consider the upper pole at $z = +L$ emitting filaments. The probability of emitting a filament into solid angle $d\Omega$ in direction $\theta$ is: $dP = f(\theta)\sin(\theta)d\theta d\phi$, where the $\sin(\theta)$ factor accounts for the spherical coordinate measure. To find the density at position $(z, r)$, we need to transform from spherical emission coordinates to cylindrical observation coordinates.

The transformation relationships are:

- Distance from pole: $R^2 = r^2 + (z - L)^2$

- Radial coordinate: $r = R\sin(\theta)$
- Axial coordinate: $z - L = -R\cos(\theta)$
- Angle: $\theta =$ atan2$\left(r, -(z - L)\right)$

The Jacobian for the transformation from $(R, \theta)$ to $(r, z)$ has determinant $|J| = 1/R$. Including the radial integration measure and transforming the probability element:

$$dP = f(\theta)\sin(\theta)d\theta d\phi dR = f(\theta)(r/R)(drdz/R)d\phi R = f(\theta)rdrdzd\phi$$

The density is defined such that the number of filaments in volume element $rdrdzd\phi$ is $\rho(r, z)rdrdzd\phi$. Therefore:

$$\rho_1(r, z) = \frac{f\left(\text{atan2}\left(r, -(z - L)\right)\right)}{r^2 + (z - L)^2}$$

By symmetry, the density from the lower pole is:

$$\rho_2(r, z) = \frac{f\left(\text{atan2}(r, z + L)\right)}{r^2 + (z + L)^2}.$$

##### *S3.4 Transport Velocity in the Continuum Limit*

When a chromosome binds to a microtubule, it moves toward the source pole with velocity directed along the filament. The unit vectors pointing from position $(z, r)$ toward each pole are:

$$\hat{\mathbf{u}}_1 = \frac{(L - z, -r)}{\sqrt{(L - z)^2 + r^2}}$$

$$\hat{\mathbf{u}}_2 = \frac{(-L - z, -r)}{\sqrt{(-L - z)^2 + r^2}}$$

In the continuum limit, where binding and unbinding occur rapidly, the system reaches a steady state where the mean velocity is the sum of contributions from motors pulling toward each pole. This velocity is proportional to the number of engaged motors and the transport velocity per motor, and depends on the steady-state binding probabilities. The explicit form of this velocity is given by the mean-field approximation in section S4.2.

##### *S3.5 Discrete Element Implementation*

In the computational model, we implement this transport mechanism using discrete force-generating elements. Each chromosome has $n$ independent elements (typically 10-20) that can be in one of three states:

- State -1: Bound to a microtubule from pole 1
- State 0: Unbound
- State +1: Bound to a microtubule from pole 2

The net velocity is the sum of contributions from all bound elements:

$$\mathbf{v} = \sum_i v_{\text{scale}}\, s_i \hat{\mathbf{u}}_{\text{pole}}(s_i)$$

where $s_i$ is the state of element $i$ and $\hat{\mathbf{u}}_{\text{pole}}$ points toward the appropriate pole.

##### *S3.6 Binding Kinetics: Saturated and Unsaturated Regimes*

The binding dynamics follow different kinetics depending on the experimental conditions:

**Unsaturated Regime (General Case):**
When motor proteins are limiting or microtubule density varies significantly, binding follows Michaelis-Menten kinetics:

$$P_{\text{bind},i} = k_{\text{on}} dt \left( \frac{\rho_i}{K_m + \rho_i} \right)$$

where $K_m$ is the Michaelis constant representing the half-saturation density. This formulation captures the dependence on binding sites (spindle microtubules).

**Saturated Regime (High Density Limit):**
When microtubule density greatly exceeds $K_m$ throughout the spindle ($\rho_i \gg K_m$), the Michaelis-Menten expression simplifies:

$$\lim_{\rho_i \gg K_m} \left( \frac{\rho_i}{K_m + \rho_i} \right) = 1$$

Therefore: $P_{\text{bind}} = k_{\text{on}} dt$ (independent of position)

This saturated regime represents control conditions where transient interactions are limited by motor kinetics rather than microtubule density. The steady-state probability of being bound to one pole when both are accessible is:

$$P_{\text{bound}} = \frac{k_{\text{on}}}{k_{\text{off}} + 2k_{\text{on}}}$$

##### *S3.7 Force Direction: Straight vs. Curved Filaments*

The direction of force depends on the geometry of microtubule trajectories. In the saturated limited, since the binding probability no longer depends on local density, we can formulate the model with other microtubule geometries.

**Straight Filaments:**
Forces point directly toward the poles along straight lines, using the unit vectors $\hat{\mathbf{u}}_1$ and $\hat{\mathbf{u}}_2$ defined earlier.

**Curved Filaments:**
Real spindle microtubules often follow curved paths. We model these using Lp-norm geometry:

$$\| \mathbf{x} \|_p = [|z|^p + |r/a|^p]^{1/p}$$

where $p = 2 +$ curvature (typically $p = 8$) and $a$ is the aspect ratio. The force direction follows the gradient of this norm:
For pole 1: $\text{term}_1 = (L - z)^p + (r/a)^p$

$$v_{z,1} = -(L - z)^{p-1} \text{term}_1^{(1/p)-1}$$

$$v_{r,1} = -r^{-1} (r/a)^p \text{term}_1^{(1/p)-1}$$

These components are then normalized to obtain unit vectors. The velocity is set to zero on the spindle axis for the radial component, while preserving axial motion.

##### *S3.8 Simulation Algorithm*

The transient model simulation uses sub-timestep integration for accurate stochastic dynamics:

1. Initialize positions and all elements to unbound state
2. For each sub-timestep $dt_{\text{sub}} = 0.01s$:
   a. Calculate current position-dependent quantities (densities $\rho_i$ for unsaturated, force directions)
   b. For each force element:
      – If bound: Check unbinding with $P_{\text{unbind}} = k_{\text{off}} dt_{\text{sub}}$
      – If unbound: Check binding to each pole with appropriate probabilities

   c. Sum forces from all bound elements
   d. Update position: $\mathbf{x}(t + dt) = \mathbf{x}(t) + \mathbf{v}dt + \sigma\sqrt{dt}\boldsymbol{\eta}$
3. Downsample to experimental time resolution

#### S4. Analytical Solutions

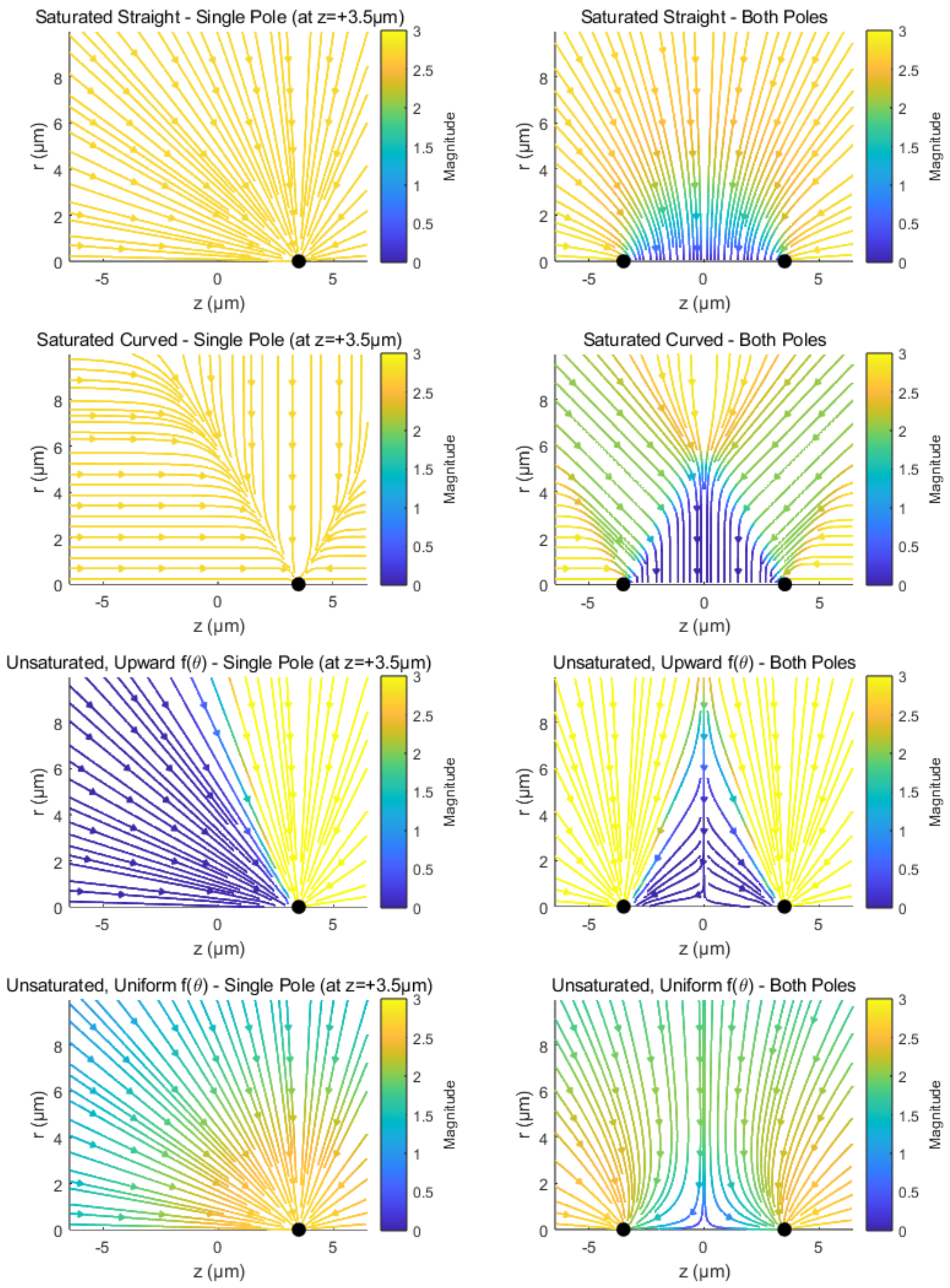

##### *S4.1 Stable Model - Analytical Velocity*

For the stable model at arbitrary position $(z, r)$, the analytical velocities are:

$$\frac{dz}{dt} = \frac{r_2 v_r(r_2) - r_1 v_r(r_1)}{2L}$$

$$\frac{dr}{dt} = \frac{r_1 v_r(r_1)(z+L) - r_2 v_r(r_2)(z-L)}{2Lr}$$

where $r_1 = \sqrt{r^2 + (z-L)^2}$ and $r_2 = \sqrt{r^2 + (z+L)^2}$, and $v_r(r)$ is the (signed) rate of radius change.

At the spindle midplane ($z = 0$), these simplify by symmetry ($r_1 = r_2$):

- $v_z = dz/dt = 0$
- $v_r = dr/dt = \frac{\sqrt{r^2+L^2}}{r} \cdot v_r\left(\sqrt{r^2 + L^2}\right)$

For the decomposed velocities at the midplane:

- Velocity toward center: $v_{\text{center}} = -v_r$ (This is positive for inward motion if $v_r$ is negative)
- Orthogonal velocity: $v_{\text{ortho}} = v_z = 0$

##### *S4.2 Transient Model - Mean-Field Approximation*

The mean-field velocity for the transient model with $n$ force elements is:

$$\mathbf{v} = n\left(P_{\text{bound},1} v_{\text{scale},1} \hat{\mathbf{u}}_1 + P_{\text{bound},2} v_{\text{scale},2} \hat{\mathbf{u}}_2\right)$$

where $P_{\text{bound},i}$ is the steady-state probability of a single force element binding to pole $i$.

**Saturated case:**

$$P_{\text{bound},1} = P_{\text{bound},2} = \frac{k_{\text{on}}}{k_{\text{off}} + 2k_{\text{on}}}$$

**Unsaturated case:**
The effective binding rate to each pole, $k_{\text{bind},i}$, follows Michaelis-Menten kinetics dependent on the local microtubule density $\rho_i$:

$$k_{\text{bind},i} = k_{\text{on}} \frac{\rho_i}{K_m + \rho_i}$$

The steady-state probability of being bound to pole $i$ is then given by:

$$P_{\text{bound},i} = \frac{k_{\text{bind},i}}{k_{\text{off}} + k_{\text{bind},1} + k_{\text{bind},2}}$$

where the densities $\rho_i$ are calculated as in S3.3.

At the midplane ($z = 0$) with symmetric poles:

- Velocity toward center: $v_{\text{center}} = -v_r$
- Orthogonal velocity: $v_{\text{ortho}} = v_z = 0$

For straight filaments at $z = 0$:

$$v_{\text{center}} = \frac{2nP_{\text{bound}}v_{\text{scale}}r}{\sqrt{r^2 + L^2}}$$

The positive sign indicates inward motion toward the spindle center.

#### S5. Parameter Values and Justification

##### *S5.1 Common Simulation Parameters*

**Table S1. Parameters common to all simulations**

| Parameter | Symbol | Value | Units | Description |
|---|---|---|---|---|
| Chromosomes per simulation | $n_{\text{chr}}$ | 920 | - | Total chromosomes (46 × 20 cells) |
| Number of cells | $n_{\text{cells}}$ | 20 | - | Simulated cell count |
| Output time points | $n_{\text{timepoints}}$ | 48 | - | Matches experimental resolution |
| Output time step | $dt$ | 5 | s | Experimental frame rate |
| Simulation timestep | $dt_{\text{sub}}$ | 0.01 | s | Internal integration step |
| Half-spindle length | $L$ | 3.5 | $\mu$m | Measured from experimental data |
| Velocity calculation window | $dTV$ | 15 | s | Window for finite differences |
| Velocity threshold | $v_{\text{thresh}}$ | 8 | $\mu$m/min | Jump artifact filter |
| Number of distance bins | $n_{\text{bins}}$ | 21 | - | For velocity profiles |

##### *S5.2 Model-Specific Parameters*

**Table S2. Stable Attachment Model parameters**

| Parameter | Symbol | Scenario 4: Constant | Scenario 5: PEF | Units | Description |
|---|---|---|---|---|---|
| Contraction function | $v_r(r)$ | -0.02 | $-0.02(1 - 3e^{-r/2})$ | $\mu$m/s | Sphere radius rate of change (negative for contraction) |
| Noise amplitude 1 | $\sigma_1$ | 0.1 | 0.1 | $\mu$m/$\sqrt{\text{s}}$ | Radius change noise |
| Noise amplitude 2 | $\sigma_2$ | 0.025 | 0.025 | $\mu$m/$\sqrt{\text{s}}$ | Position noise |
| PEF decay length | $r_0$ | - | 2 | $\mu$m | Polar ejection force scale |
| PEF strength | $\alpha$ | - | 3 | - | Maximum PEF effect |

**Table S3. Transient Model parameters**

| Parameter | Symbol | Saturated (Scenarios 1,3) | Unsaturated (Scenario 2) | Units | Description |
|---|---|---|---|---|---|
| Binding rate | $k_{\text{on}}$ | 10 | 10 | $s^{-1}$ | Maximum binding rate |
| Unbinding rate | $k_{\text{off}}$ | 1 | 1 | $s^{-1}$ | Detachment rate |
| Velocity scale | $v_{\text{scale}}$ | 0.005 | 0.005 | $\mu$m/s | Base transport velocity |
| Force elements | $n_{\text{int}}$ | 10 | 20 | - | Per chromosome |
| Michaelis constant | $K_m$ | 0 (saturated limit) | $10^{-3.5}$ | $\mu m^{-2}$ | Half-saturation density |
| Curvature | $p-2$ | 6 (S1), 0 (S3) | 0 (straight) | - | Filament path curvature |
| Aspect ratio | $a$ | 1.5 (S1), 1 (S3) | 1 | - | Spindle shape factor |
| Angular distribution | $f(\theta)$ | - | See Eq. S6.1 | - | MT angle preference |
| Distribution params | $\kappa, \mu$ | - | 7.5, $\pi/2$ | -, rad | Angular sigmoid parameters |

##### *S5.3 Angular Distribution Function*

The angular sigmoid distribution used for the unsaturated model is:

$$f(\theta) = \frac{\kappa(1 + \tanh[\kappa(\theta - \mu)])}{2\pi\kappa + \log\left[\cosh\left(\kappa(2\pi - \mu)\right) \cdot \text{sech}(\kappa\mu)\right]} \quad \text{(Eq. S5.1)}$$

where $\kappa = 7.5$ controls the distribution width and $\mu = \pi/2$ sets the threshold angle. This form provides smooth variation from uniform ($\kappa \to 0$) to strongly peaked ($\kappa \gg 1$) distributions for values above $\theta = \mu$, representing the preferential emission of microtubules perpendicular to the spindle axis under certain conditions.

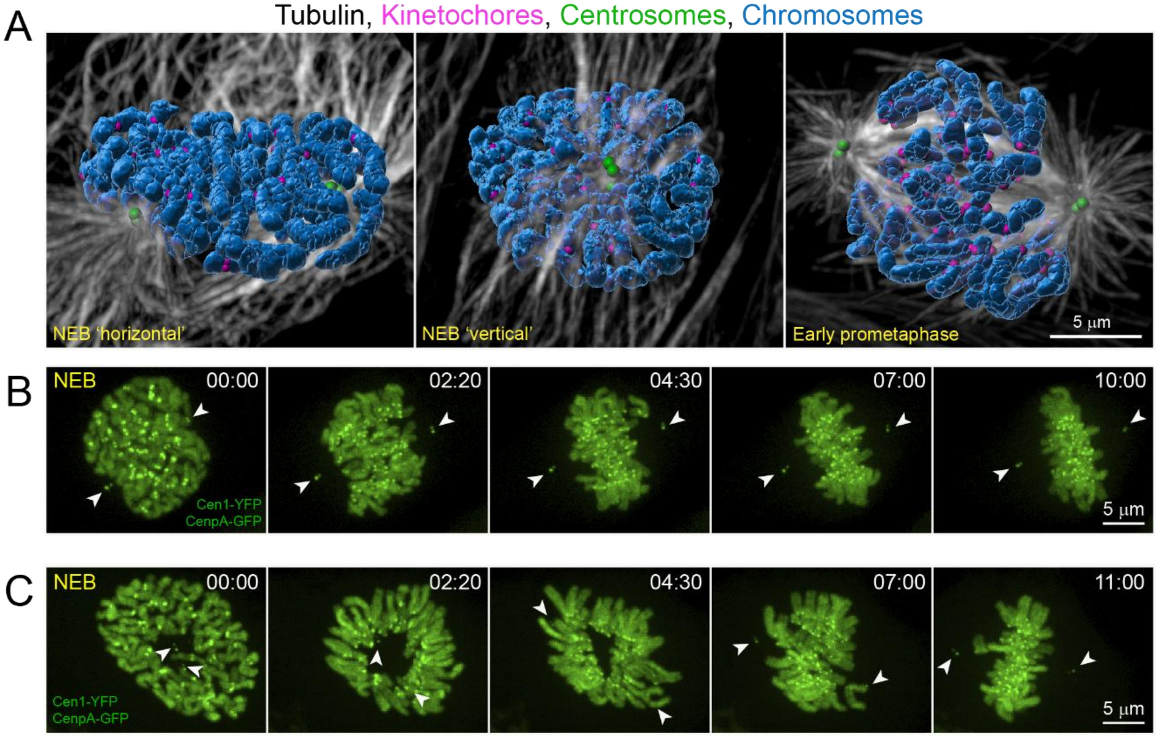


**Fig. S1. Spatial distributions of chromosomes during prometaphase**. (A) 3D-surface renderings of fixed RPE1 cells that express centrin1-GFP (centrosomes), CenpA-GFP (kinetochores), and immunostained for α-Tubulin (microtubules). Chromosomes are counterstained with Hoechst 33342, segmented, and surface rendered. At nuclear envelope breakdown (NEB), duplicated centrosomes reside within invaginations of the nuclear envelope on the opposite sides of the nucleus, either along its longer (NEB 'horizontal') or the shorter (NEB 'vertical') axis. In early-mid prometaphase chromosomes are not present in the immediate proximity of the centrosomes/spindle poles. (B-C) Selected time points from 4-D recordings of RPE1 cells with centrosomes separated along the longer (B) or shorter (C) axis of the nucleus at NEB. Maximum-intensity projections of the entire cell. 3D views of these cells are shown in Figure 1 C-D. Arrowheads mark centrosomes/spindle poles. Time in minutes : seconds from NEB.

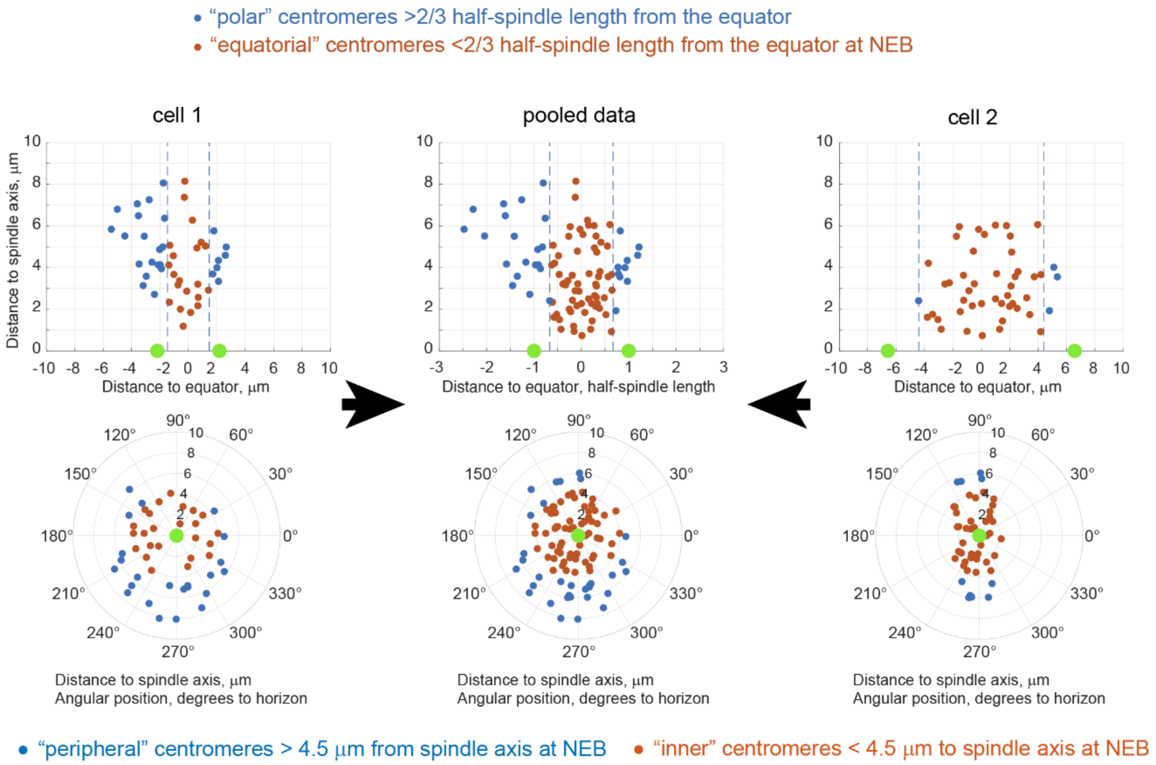


**Fig. S2. Approach for quantification of centromere behavior.** Centromere positions are expressed in a cylindrical coordinate system with the origin at the center of the spindle. Plots present the distance to the spindle axis ($\rho$) vs. the distance to the equatorial plane ($Z$) or the distance to the spindle axis ($\rho$) vs. the angular position ($\theta$). To discriminate ‘bioriented’ ($Z$ < 2/3 of the half-spindle length from the equator) vs. ‘monooriented’ ($Z$ > 2/3 of the half-spindle length from the equator) chromosomes in datasets containing multiple cells, $Z$ values are normalized to the half of the spindle length at each timepoint.

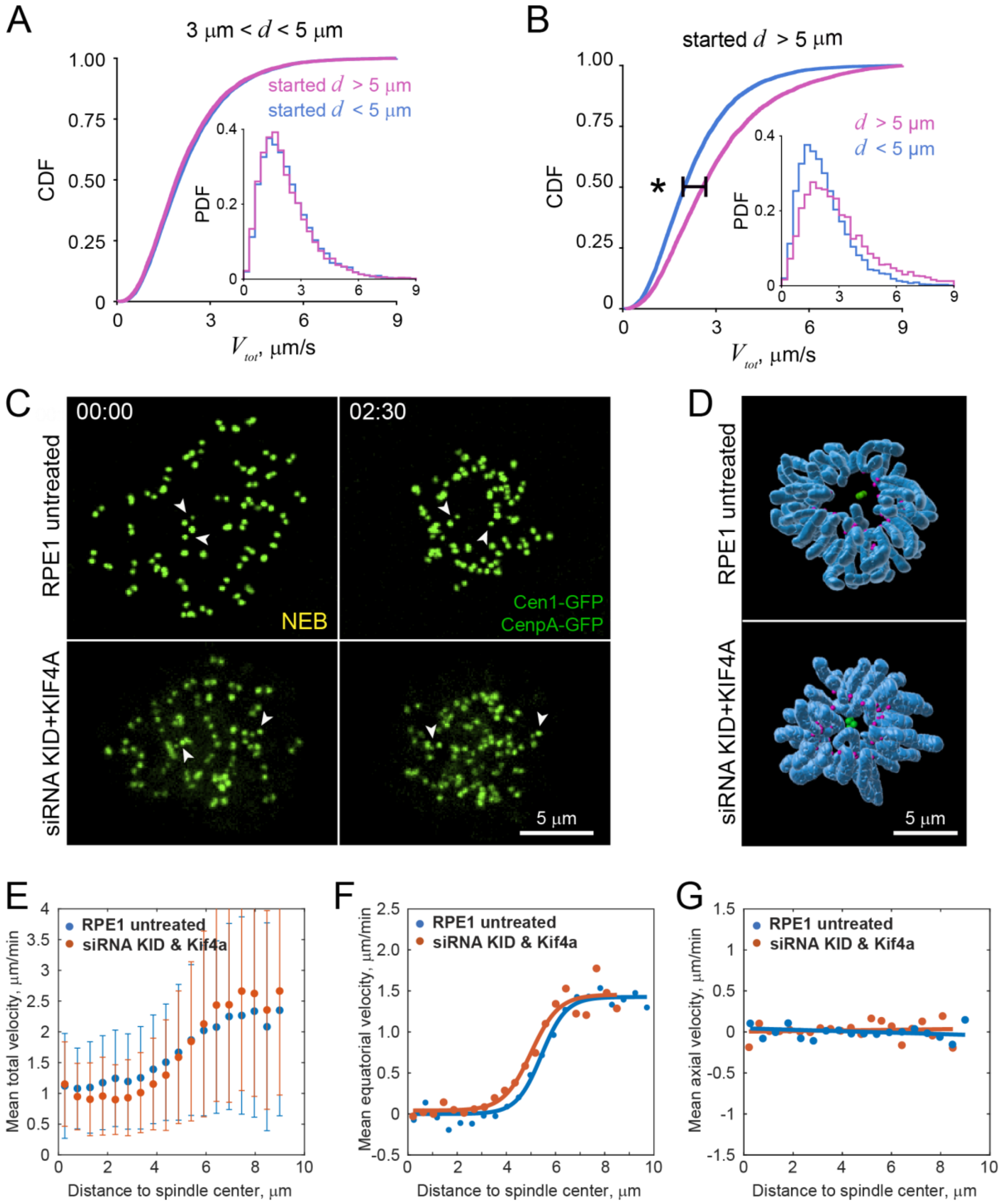


**Fig. S3. Slower centromere movements are not caused by crowding within the inner parts of the spindle neither by the spindle ejection forces. (A)** Cumulative distribution function (CDF) and probability density function (PDF) of velocities for centromeres initially positioned far (>5 μm, magenta) vs. closer (<5 μm, blue) from the spindle center when they move through the intermediate region of the spindle (3-5 μm from the center). No significant difference exists (two-sample $t$-test, $T$ = 1.8912 and $df$ = 17958). **(B)** CDF and PDF of velocities for centromeres that are initially positioned >5-μm away from the spindle center when they move through the peripheral (>5 μm from the center) vs. inner (<5 μm from the center) parts of the spindle center. Significant different (two-sample $t$-test, $T$ = -32.8887 and $df$ = 16812). **(C-D)** Effect of siRNA co-depletion of chromokinesins Kid and Kif4a on chromosome distribution and orientation during mid-prometaphase. Notice the presence of kinetochores near the spindle axis in the chromokinesins-depleted cell. This zone is reproducibly devoid of kinetochores in the untreated RPE1. (C) Maximum-intensity projections of the entire cell from time-lapse recordings of cells with GFP-tagged kinetochores and centrioles. Arrowheads denote centrioles. (D) Surface renderings of chromosome arms (blue) with marked positions of kinetochores (magenta) and centrioles (green). **(E)** Centromere total velocity as a function of distance to the spindle center in untreated RPE1 vs. RPE1 co-depleted for Kid and Kif4a. No significant change is observed. **(F-G)** similar to (E) but only the equatorial (F) and axial (G) components of centromere velocity are plotted. No significant differences are observed.

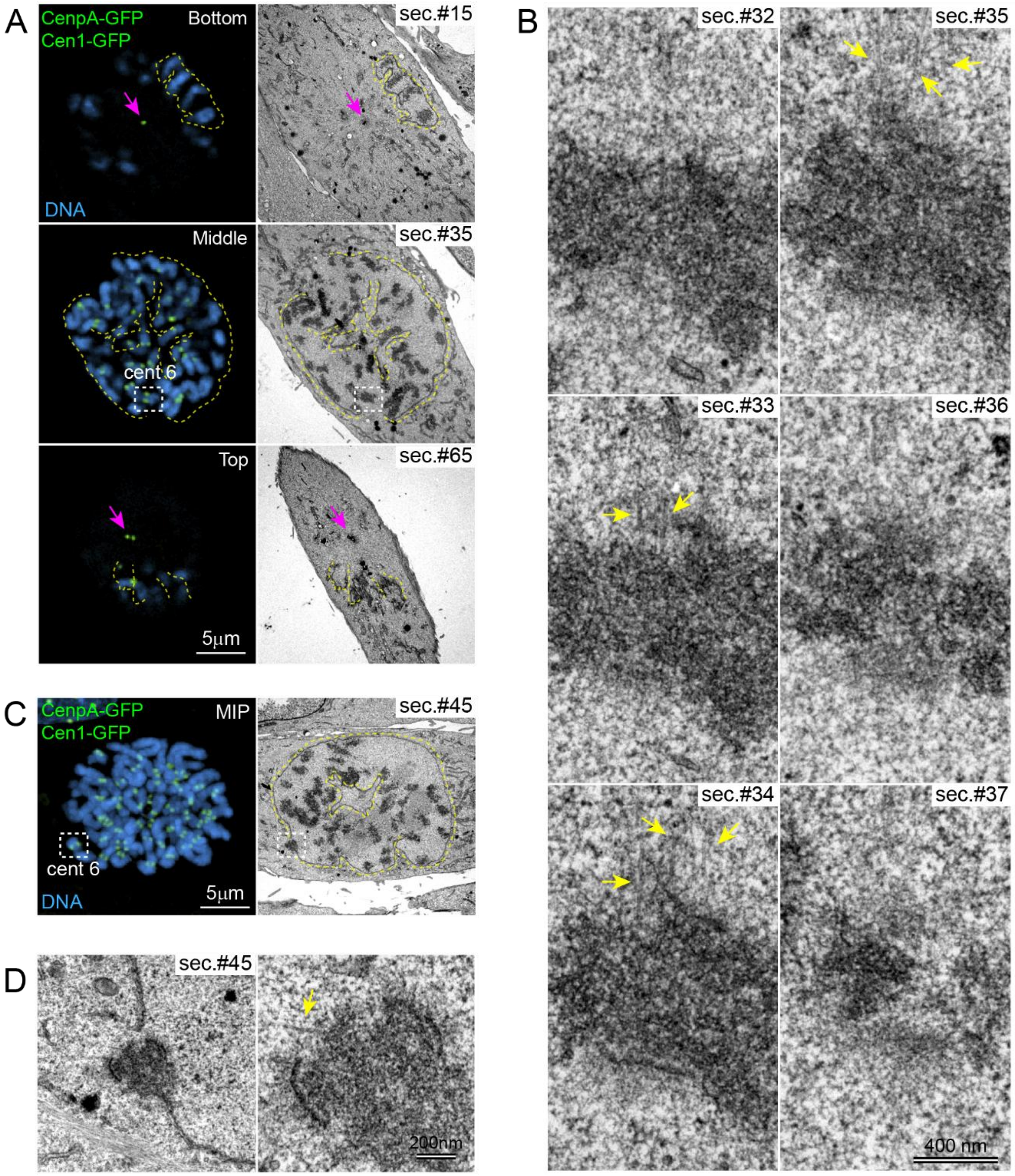


**Fig. S4. Correlative Light/Electron Microscopy (CLEM) analysis of RPE1 cells fixed during nuclear envelope breakdown. (A)** Selected LM planes and corresponding 80-nm EM sections of an RPE1 cell with fenestrated nuclear envelope (3-D reconstruction of this cell shown in Figure 5B). Centromeres and centrioles are tagged with CenpA-GFP and Centrin1-GFP. Chromosomes are stained with Hoechst 33342. Arrows mark centrioles, located within deep invaginations of nuclear envelope on the ventral and dorsal sides of the nucleus. Dashed yellow lines denote nuclear envelope (traced in EM sections). Discontinuities in the nuclear envelope (fenestrae) are evident in EM sections through the medial part of the cell (sec.#35). Note that the fenestrae are not adjacent to the centrosomes. Area within the dashed white box is shown in (B). (B) Serial 80-nm sections through the centromere of one chromosome (boxed in A). Arrows point at microtubules emanating from one kinetochore. No microtubule is present at the sister kinetochore. Section #34 and traces of microtubules are also shown in Figure 5C. (C) Similar to (A) but this cell was fixed at the earlier stage with just two small fenestrae forming near a small chromosome. **(D)** Higher magnifications of the centromere (cent6) adjacent to a fenestra (boxed in C) reveal a single short microtubule (arrow) near the centromere.

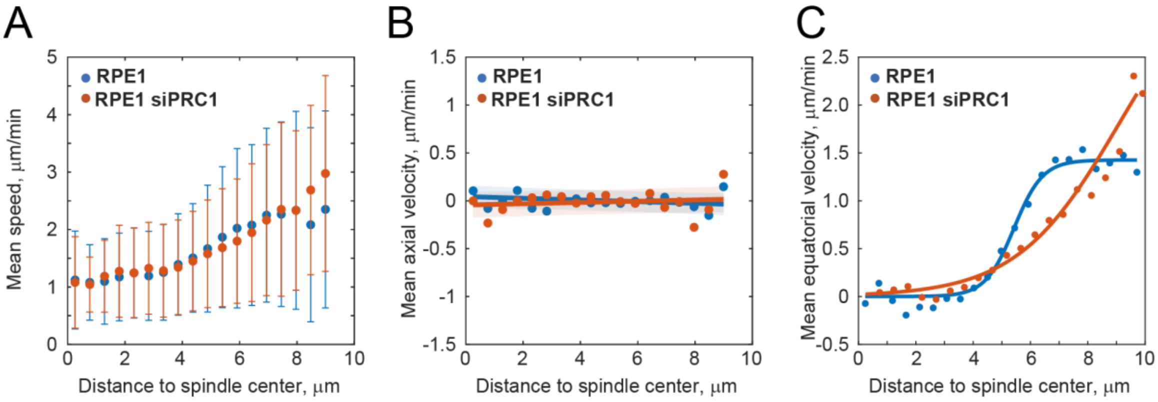


**Fig. S5. Abrogation of microtubule bundling within the spindle changes relationship between centromere velocity and distance to the spindle center. (A)** Centromere total velocity as a function of distance to the spindle center in untreated RPE1 vs. RPE1 depleted for PRC1 via shRNA. **(B-C)** similar to (A) but only the axial (B) and equatorial (C) components of centromere velocity are shown. Notice that equatorial velocity increases slower in between 4 and 8 μm and continues to increase at larger distances from the spindle center.
